# Kinetochore recruitment of CENP-F illustrates how paralog divergence shapes kinetochore composition and function

**DOI:** 10.1101/276204

**Authors:** Giuseppe Ciossani, Katharina Overlack, Arsen Petrovic, Pim Huis in ‘t Veld, Carolin Körner, Sabine Wohlgemuth, Stefano Maffini, Andrea Musacchio

## Abstract

The metazoan proteins CENP-E and CENP-F are components of a fibrous layer of mitotic kinetochores named the corona. Several features suggest that CENP-E and CENP-F are paralogs: they are very large (approximately 2700 and 3200 residues, respectively), rich in predicted coiled-coil structure, C-terminally prenylated, and endowed with microtubule-binding sites at their termini. In addition, CENP-E contains an ATP-hydrolyzing motor domain that promotes microtubule plus-end directed motion. Here, we show that CENP-E and CENP- F are recruited to mitotic kinetochores independently of the Rod-Zwilch-ZW10 (RZZ) complex, the main corona constituent. We identify selective interactions of CENP-E and CENP-F respectively with BubR1 and Bub1, paralogous proteins involved in mitotic checkpoint control and chromosome alignment. While BubR1 is dispensable for kinetochore localization of CENP-E, Bub1 is stringently required for CENP-F localization. Through biochemical reconstitution, we demonstrate that the CENP-E:BubR1 and CENP-F:Bub1 interactions are direct and require similar determinants, a dimeric coiled-coil in CENP-E or CENP-F and a kinase domain in BubR1 or Bub1. Our findings are consistent with the existence of ‘pseudo-symmetric’, paralogous Bub1:CENP-F and BubR1:CENP-E axes, supporting evolutionary relatedness of CENP-E and CENP-F.

## Introduction

The segregation of chromosomes from a mother cell to its daughters during cell division relies on the function of specialized protein complexes, the kinetochores, as bridges linking chromosomes to spindle microtubules (Musacchio and Desai, 2017). Kinetochores are built on specialized chromosome loci known as centromeres, whose hallmark is the enrichment of the histone H3 variant centromeric protein A (CENP-A, also known as CenH3) (Earnshaw, 2015). CENP-A seeds kinetochore assembly by recruiting CENP-C, CENP-N, and their associated protein subunits in the constitutive centromere associated network (CCAN) (Cheeseman and Desai, 2008). These centromere proximal ‘inner kinetochore’ subunits, in turn, recruit the centromere distal ‘outer kinetochore’ subunits of the KMN complex (<u>K</u>nl1 complex, <u>M</u>is12 complex, <u>N</u>dc80 complex), which promote ‘end-on’ microtubule binding and control the spindle assembly checkpoint (SAC) (Musacchio and Desai, 2017).

Early in mitosis, prior to end-on microtubule attachment, an additional fibrous structure, the kinetochore corona, assembles as the outermost layer of the kinetochore (Figure 1A) (Jokelainen, 1967; Magidson et al., 2015; McEwen et al., 1993; Rieder, 1982). The corona’s main constituent is a trimeric protein complex named RZZ [from the name of the fruit fly genes Rough Deal (ROD), Zwilch, and Zeste White 10 (ZW10)]. The ROD subunit is structurally related to proteins that oligomerize near biological membranes to promote vesicular trafficking, including Clathrin (Civril et al., 2010; Mosalaganti et al., 2017), leading to hypothesize that corona assembly results from RZZ polymerization (Mosalaganti et al., 2017). The interaction of the RZZ complex with an adaptor subunit named Spindly, in turn, further recruits the microtubule minus-end directed motor cytoplasmic Dynein and its binding partner Dynactin to kinetochores, as well as the Mad1:Mad2 complex, which is crucially required for SAC signaling (Barisic et al., 2010; Basto et al., 2004; Buffin et al., 2005; Caldas et al., 2015; Chan et al., 2009; Cheerambathur et al., 2013; Gama et al., 2017; Gassmann et al., 2008; Gassmann et al., 2010; Griffis et al., 2007; Howell et al., 2001; Kops et al., 2005; Matson and Stukenberg, 2014; Mische et al., 2008; Silio et al., 2015; Sivaram et al., 2009; Starr et al., 1998; Varma et al., 2008; Williams et al., 1996; Wojcik et al., 2001; Yamamoto et al., 2008; Zhang et al., 2015).

**Figure 1.**
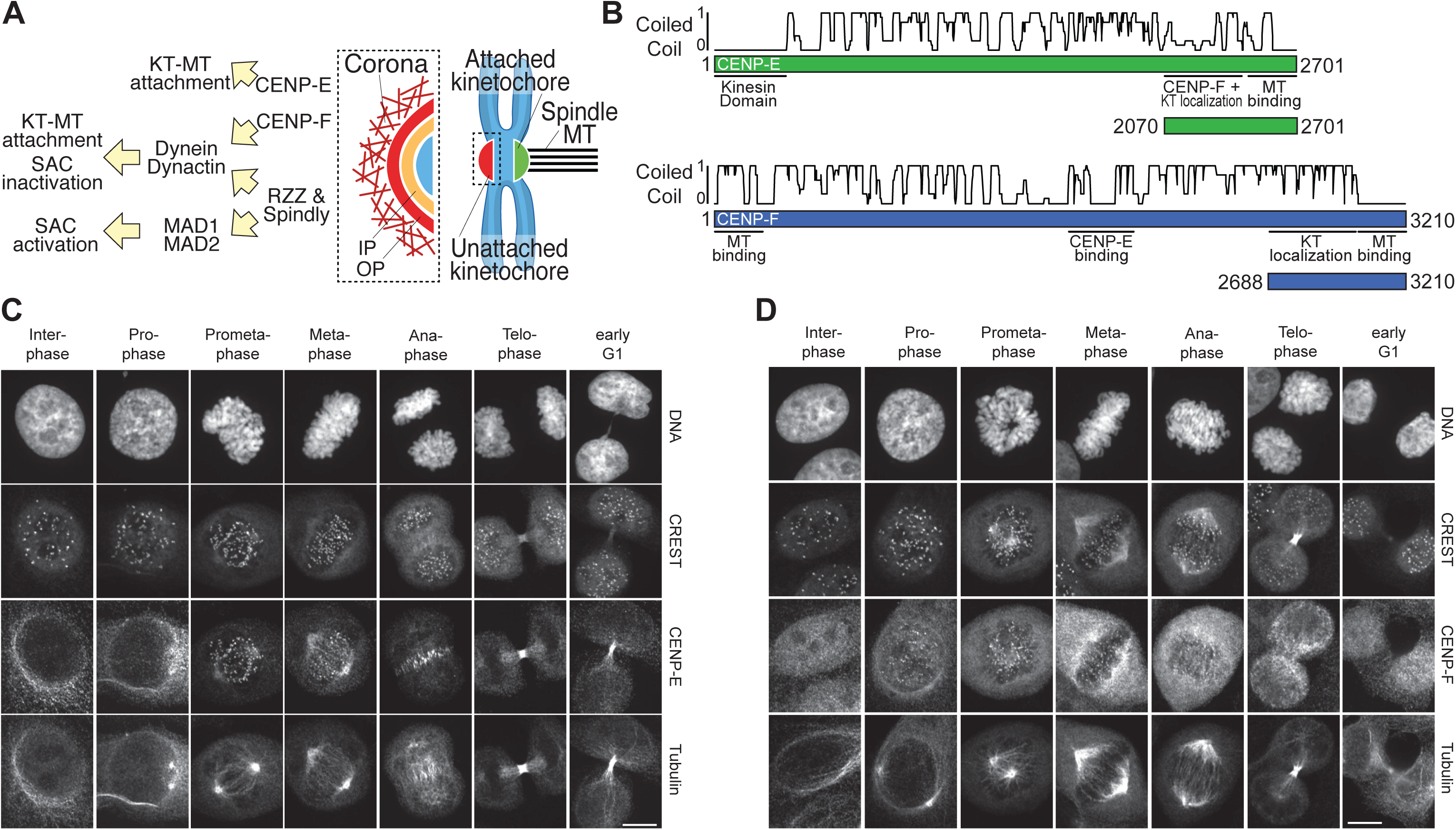
The corona proteins CENP-E and CENP-F localize at kinetochores with distinct timing. **A**) Schematic representation of the corona structure and function. MT, microtubules; IP, inner plate of kinetochore; OP, outer plate of kinetochore. **B**) Schematic organization of the CENP-E and CENP-F full-length proteins. Coiled coil regions were predicted with coils, pcoils and marcoils (Delorenzi and Speed, 2002; Gruber et al., 2005; Lupas et al., 1991) using default parameters (ncoils and paircoils: windows size 21). To combine all three coiled coil prediction algorithms, we applied a scoring system in which we assigned for each residue two points for a high significance (P-value >=0.9) and one point for low significance (P-value >=0.8). Two additional points were granted for an identical register position in coiled coil if predicted by all three programs, resulting in a maximum score of 8. **C, D**) Representative images of fixed Hela cells treated for fluorescence staining with the indicated antibodies. The panel illustrates the localization of CENP-E (C) and CENP-F (D) in the different phases of the cell cycle. Scale bar: 10 µm.

**Figure 1S1.**
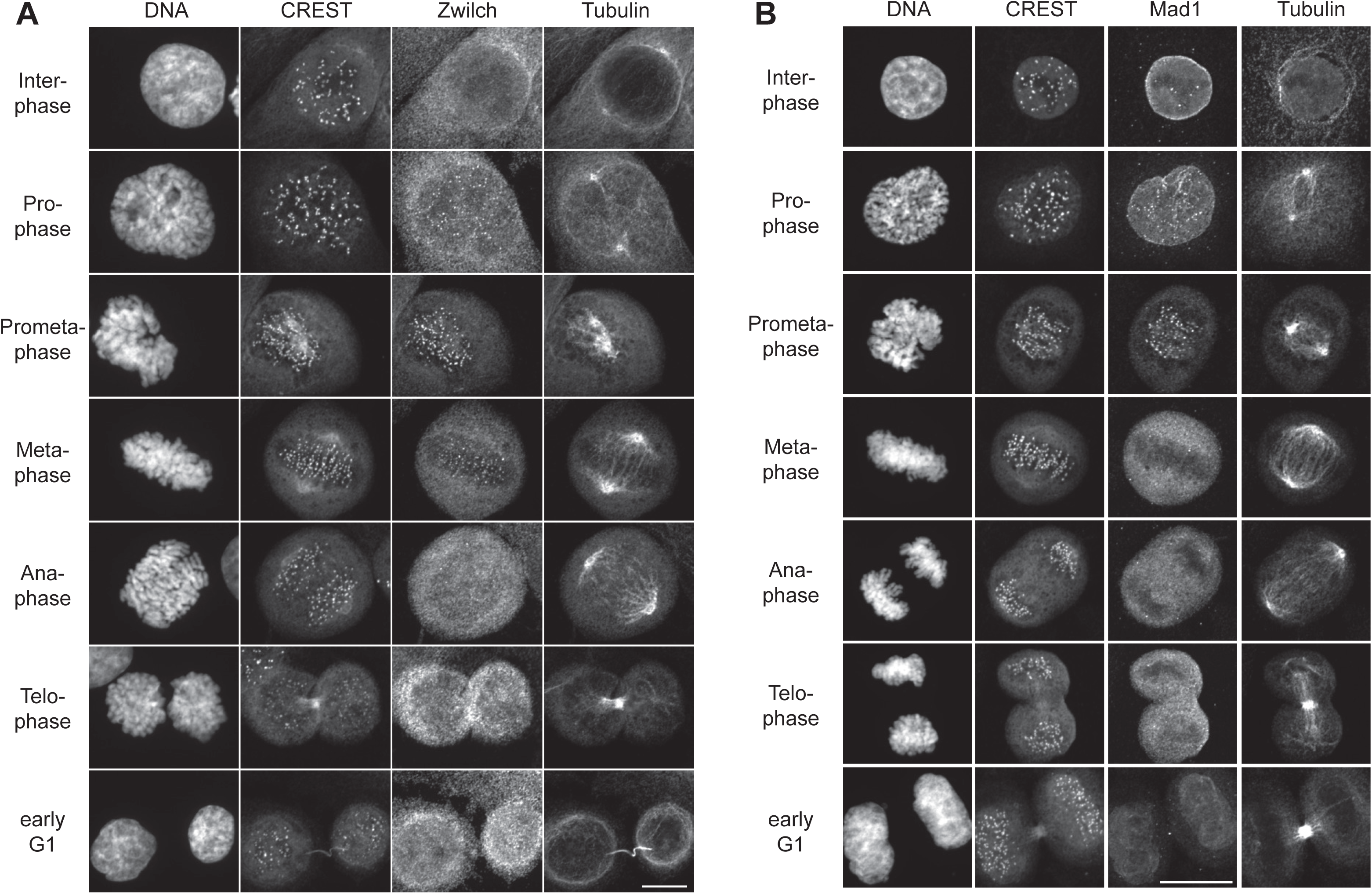
Zwilch and Mad1 localization through the cell cycle. **A**, **B**) Representative images of Hela cells showing Zwilch (A) and Mad1 (B) localization in the different phases of the cell cycle. Scale bar: 10 µm.

Corona assembly leads to a broad expansion of the microtubule-binding interface of kinetochores that may promote initial microtubule capture, congression towards the metaphase plate, and SAC signaling (Basto et al., 2000; Buffin et al., 2005; Hoffman et al., 2001; Kops et al., 2005; Magidson et al., 2011; Magidson et al., 2015; Wynne and Funabiki, 2015). Differently from the mature end-on attachments, initial attachments of kinetochores engage the microtubule lattice and are therefore defined as lateral or side-on. CENP-E, a kinesin-7 family member, plays a crucial role at this stage. Its inhibition or depletion lead to severe and persistent chromosome alignment defects, with numerous chromosomes failing to congress towards the spindle equator and stationing near the spindle poles, causing chronic activation of the SAC (Kapoor et al., 2006; Kuhn and Dumont, 2017; Magidson et al., 2011; Magidson et al., 2015; Putkey et al., 2002; Schaar et al., 1997; Wood et al., 1997; Yao et al., 2000; Yen et al., 1991). Human CENP-E consists of 2701 residues (Figure 1B) (Yen et al., 1992). Besides the globular N-terminal motor domain, the rest of the CENP-E sequence forms a flexible and highly elongated (∼230 nm) coiled-coil (Kim et al., 2008). The kinetochore-targeting domain of CENP-Eencompasses residues 2126-2476 and is followed by a microtubule-binding region (Figure 1B) (Chan et al., 1998; Liao et al., 1994). The distribution of CENP-E in an outer kinetochore crescent shape similar to that of the RZZ supports the notion that CENP-E is part of the kinetochore corona, but its persistence at kinetochores after disappearance of the corona suggests a corona-independent localization mechanism (Cooke et al., 1997; Hoffman et al., 2001; Magidson et al., 2015; Wynne and Funabiki, 2015; Yao et al., 1997; Yen et al., 1992).

CENP-F (also known as Mitosin, 3210 residues in humans) is also a kinetochore corona constituent during early mitosis that persists at kinetochores after corona shedding (Casiano et al., 1993; Hussein and Taylor, 2002; Liao et al., 1995; Rattner et al., 1993; Zhu, 1999; Zhu et al., 1995a; Zhu et al., 1995b). Like CENP-E, CENP-F is also highly enriched in predicted coiled-coil domains (Figure 1B), but lacks an N-terminal motor domain. Rather, it contains two highly basic microtubule-binding domains in the N-terminal 385 residues and in the C-terminal 187 residues (Feng et al., 2006; Musinipally et al., 2013; Volkov et al., 2015). Similarly to CENP-E, the kinetochore recruitment domain of CENP-F is positioned in proximity of the C-terminus [encompassing residues 2581-3210, the minimal domain tested for this function to date (Hussein and Taylor, 2002; Zhu, 1999; Zhu et al., 1995a)]. The apparent similarity of CENP-E and CENP-F extends to the fact that they are both post-translationally modified with a farnesyl prenol lipid chain (isoprenoid) on canonical motifs positioned in their C-termini (Ashar et al., 2000). These modifications contribute to kinetochore recruitment of CENP-E and CENP-F, albeit to extents that differ in various reports (Holland et al., 2015; Hussein and Taylor, 2002; Moudgil et al., 2015; Schafer-Hales et al., 2007).

Previous studies identified CENP-F and BubR1 as binding partners of CENP-E (Chan et al., 1998; Mao et al., 2003; Yao et al., 2000). BubR1 is a crucial constituent of the SAC, a molecular network required to prevent premature mitotic exit (anaphase) in cells retaining unattached or improperly attached kinetochores (Musacchio, 2015). BubR1 is a subunit of the mitotic checkpoint complex (MCC), the SAC effector (Sudakin et al., 2001). Its structure is a constellation of domains and interaction motifs required to mediate binding to other SAC proteins, and terminates in a kinase domain (Musacchio, 2015). It has been proposed that CENP-E stimulates BubR1 activity, and that microtubule capture silences it (Mao et al., 2003; Mao et al., 2005). Later studies, however, identified BubR1 as an inactive pseudokinase (Breit et al., 2015; Suijkerbuijk et al., 2012), and therefore the significance of CENP-E microtubule binding for the role of BubR1 in the SAC remains unclear. Depletion or inactivation of CENP-E, however, is compatible with a robust mitotic arrest (Schaar et al., 1997; Yen et al., 1991).

A yeast 2-hybrid (Y2H) interaction of CENP-F and Bub1 has also been reported but never validated experimentally (Chan et al., 1998). Bub1, a paralog of BubR1, retained genuine kinase activity in humans and it plays a function at the interface of mitotic checkpoint signaling and kinetochore microtubule attachment (Raaijmakers et al., 2018; Suijkerbuijk et al., 2012). Suggesting that the interaction of Bub1 and CENP-F is functionally important, previous studies identified Bub1 as being essential for kinetochore recruitment of CENP-F (Johnson et al., 2004; Klebig et al., 2009; Liu et al., 2006; Raaijmakers et al., 2018).

In our previous studies, we characterized in molecular detail how sequence divergence impacted the protein interaction potential of the human Bub1 and BubR1 paralogs (Overlack et al., 2017; Overlack et al., 2015). We described a molecular mechanism that explains how Bub1, through an interaction with a phospho-aminoacid adaptor named Bub3, can interact with kinetochores and promote the recruitment of BubR1 via a pseudo-dimeric interface (Overlack et al., 2017; Overlack et al., 2015; Primorac et al., 2013). In view of these previous studies, here we have dissected the molecular basis of the interactions of BubR1 and Bub1 with CENP-E and CENP-F. We provide strong evidence for the sub-functionalization of these paralogous protein pairs.

## Results and Discussion

### Independent kinetochore localization of CENP-E, CENP-F, and the RZZ

Using specific antibodies (see Methods), we assessed the timing and specificity of kinetochore localization of CENP-E, CENP-F, Zwilch, and Mad1. CENP-E showed perinuclear localization until prometaphase, when it first appeared at kinetochores. It persisted there until metaphase, and was then found at the spindle midzone after anaphase onset (Figure 1C). This localization, which corresponds to previous descriptions (Yen et al., 1991; Yen et al., 1992), is reminiscent of that of chromosome passenger proteins (Earnshaw and Bernat, 1991). CENP-F, on the other hand, localized to kinetochores already in prophase, where it was also temporarily visible at the nuclear envelope, and persisted there until anaphase, with progressive weakening and dispersion (Figure 1D), as noted previously (Baffet et al., 2015; Bolhy et al., 2011; Hu et al., 2013; Liao et al., 1995; Rattner et al., 1993). Also Zwilch (a subunit of the RZZ complex) and Mad1 were already visible at kinetochores in prophase, but they became invisible at these structures upon achievement of metaphase (Figure 1S1A-B), in agreement with the notion that the corona becomes dissolved upon microtubule attachment (see Introduction).

Thus, both CENP-E and CENP-F continue to localize to kinetochores well beyond the timing of removal of the RZZ complex and Mad1, suggesting that they can be retained at kinetochores independent of the corona. To test this directly, we identified conditions for optimal depletion of Zwilch, CENP-E, or CENP-F by RNA interference (RNAi) (Figure 2S1A-J). Depletion of Zwilch resulted in depletion of Mad1 from kinetochores (Figure 2S1H-J), but left the kinetochore levels of CENP-E essentially untouched (Figure 2A). This observation is in agreement with previous studies showing that Mad2, whose kinetochore localization requires Mad1 (Martin-Lluesma et al., 2002; Sharp-Baker and Chen, 2001), is also depleted from kinetochores upon depletion of other RZZ subunits (Caldas et al., 2015; Gassmann et al., 2008; Raaijmakers et al., 2018). The observation that CENP-E retains kinetochore localization under conditions in which Mad1 appears to become depleted seems inconsistent with a recent report proposing that Mad1 is required for kinetochore localization of CENP-E (Akera et al., 2015), but agrees with previous reports that failed to detect consequences on CENP-E localization upon depletion of Mad1 (Martin-Lluesma et al., 2002; Sharp-Baker and Chen, 2001).

**Figure 2.**
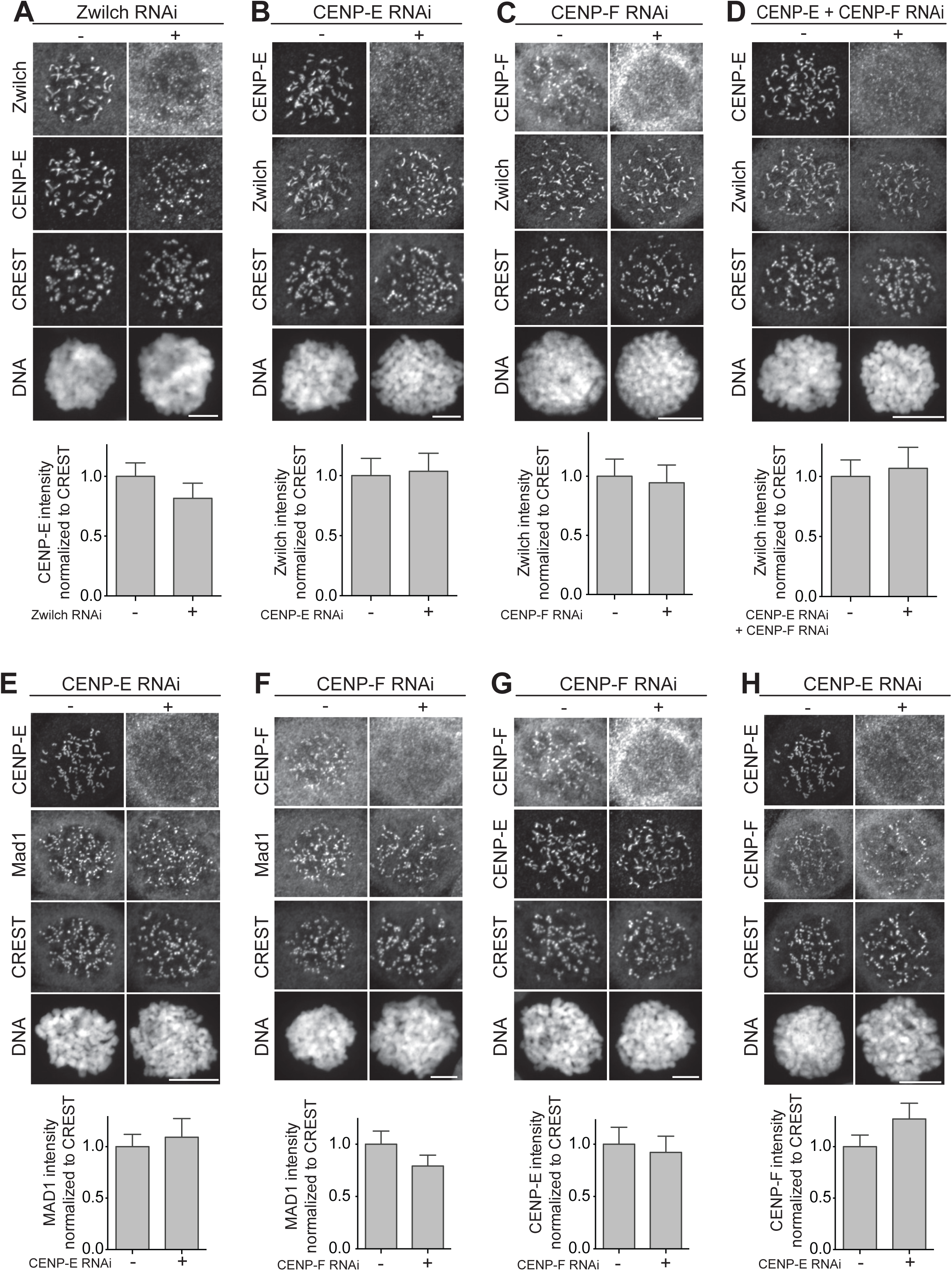
Kinetochore localization of RZZ and Mad1 are independent of CENP-E and CENP-F. **A-H**) Representative images and quantification of proteins kinetochore levels in Hela cells mock treated or depleted of Zwilch (A), CENP-E (B, E, H), CENP-F (C, F, G) or co-depleted of CENP-E and CENP-F (D). Scale bar: 10 µm. Zwilch depletion does not affect the localization of CENP-E (A). CENP-E depletion does not affect the localization of Zwilch (B), Mad1 (E) and CENP-F (H). Similarly, CENP-F depletion does not interfere with the recruitment of Zwilch (C), Mad1 (F) and CENP-E (G). Co- depletion of CENP-E and CENP-F has no effects on localization of Zwilch (D). The graphs show mean intensity of one (B, C, E), two (D, F) or three (A, G, H) experiments; the error bars indicate SEM and the mean values for non-depleted cells are set to 1.

**Figure 2S1.**
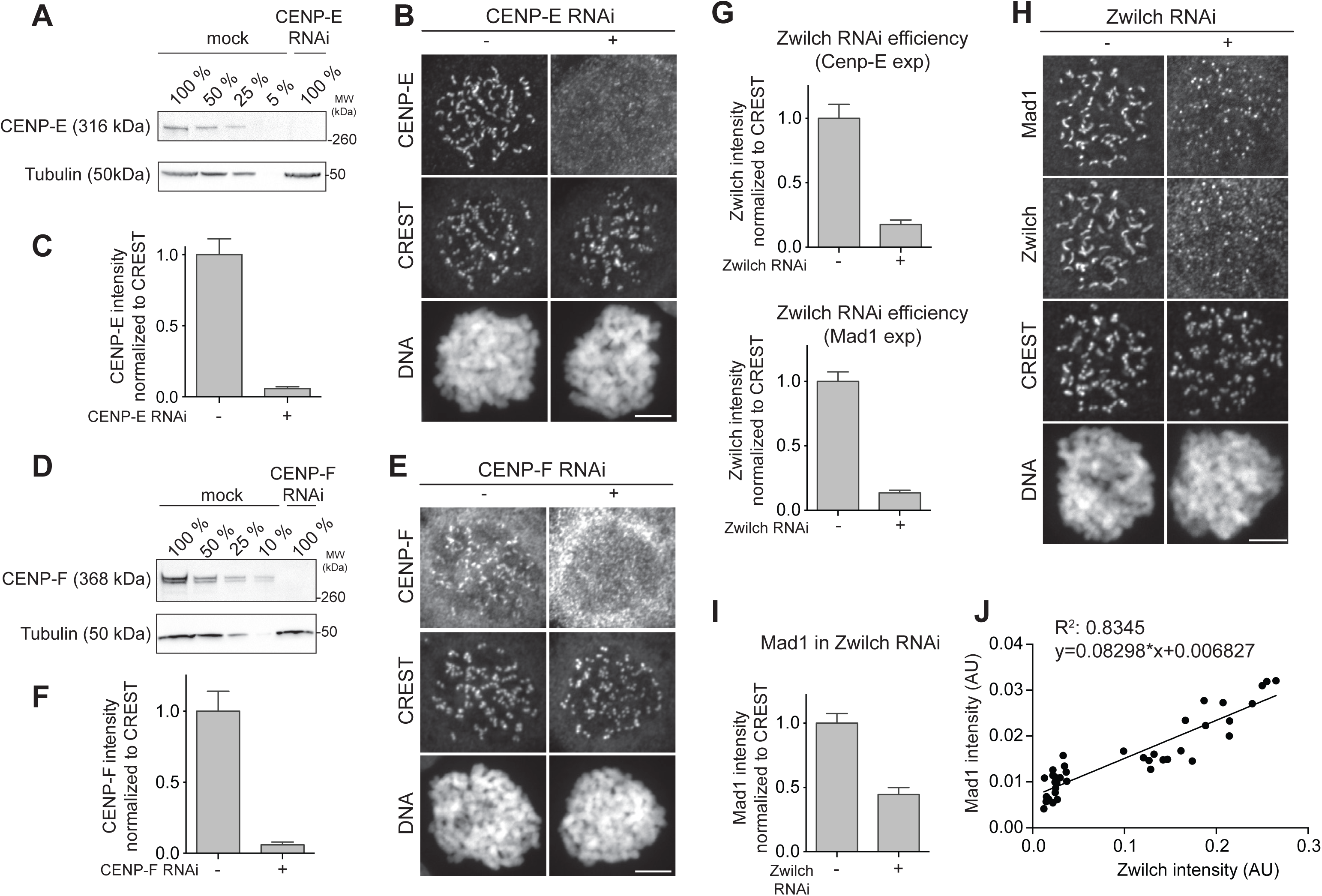
Efficient siRNA depletion of CENP-E, CENP-F and Zwilch. **A**) Western blot showing CENP-E protein levels in mock treated or CENP-E depleted cells. **B**) Representative images of HeLa cells, either mock-treated or depleted of CENP- E, showing that CENP-E can be efficiently depleted. Scale bar: 10 µm. **C**) Quantification of CENP-E kinetochore levels in cells treated as in panel (B). The graph shows mean intensity of two independent experiments; the error bars indicate SEM and the mean value for non-depleted cells is set to 1. **D**) Western blot showing CENP-F protein levels in mock treated or CENP-F depleted cells. **E**) Representative images of HeLa cells mock treated or depleted of CENP-F showing successful CENP-F depletion. Scale bar: 10 µm. **F**) Quantification of CENP-F kinetochore levels in cells treated as in panel (E). The graph shows mean intensity of three independent experiments; the error bars indicate SEM and the mean value for non-depleted cells is set to 1. **G**) Quantification of Zwilch kinetochore levels in cells treated as in Figure 2A (upper panel) and in (H). The graphs show mean intensity of three (CENP-E experiment) or two (Mad1 experiment) independent experiments; the error bars indicate SEM and the mean value for non- depleted cells is set to 1. **H**) Representative images of HeLa cells mock treated or depleted of Zwilch showing that Zwilch depletion leads to reduced Mad1 levels. Scale bar: 10 µm. **I**) Quantification of Mad1 kinetochore levels in cells treated as in (H). The graph shows mean intensity of two independent experiments; the error bars indicate SEM and the mean value for non-depleted cells is set to 1. **J**) Correlation of Mad1 and Zwilch levels in 38 cells, 19 of which were mock treated, while the other 19 were RNAi depleted of Zwilch. Cells are from two independent experiments. AU, arbitrary units.

**Figure 2S2.**
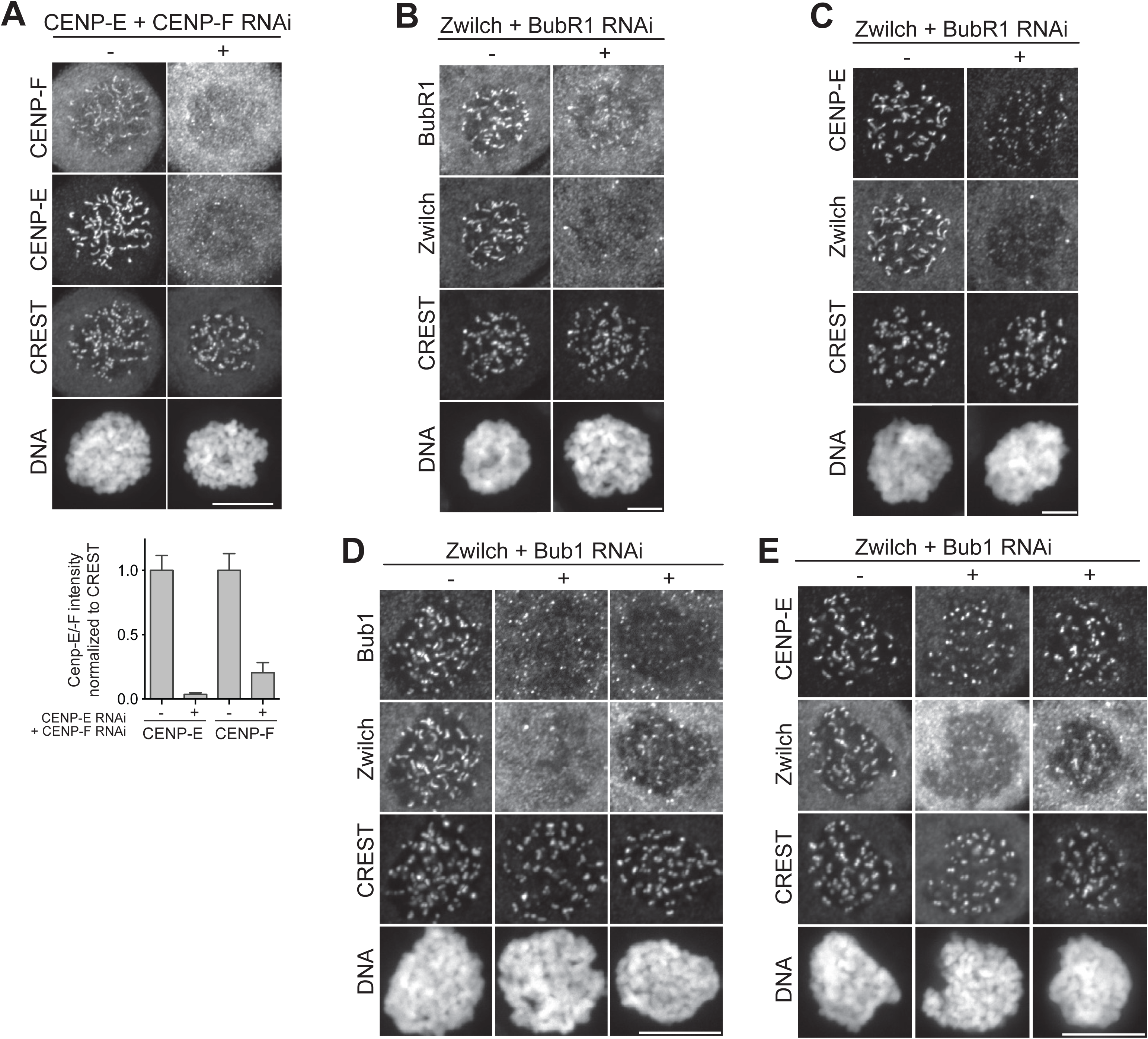
Zwilch and BubR1 co-depletion does not affect CENP-E kinetochore recruitment. **A**) Upper panel - Representative images of HeLa cells either mock treated or co-depleted of CENP-E and CENP-F showing effective co-depletion of the proteins. Scale bar: 10µm. Lower panel - Quantification of co-depletion efficiency. The graph shows mean intensity of two independent experiments; the error bars indicate SEM and the mean values for non-depleted cells are set to 1. **B**) Representative images of HeLa cells mock treated or co-depleted of Zwilch and BubR1 showing efficient depletion of both the proteins. Scale bar: 10 µm. **C**) Representative images of HeLa cells mock treated or co- depleted of Zwilch and BubR1 showing that CENP-E localization is not affected. Scale bar: 10 µm. **D**) Representative images of HeLa cells mock treated or co-depleted of Zwilch and Bub1 showing efficient depletion of both proteins. Two cells with different Zwilch levels and the same low Bub1 levels are shown for the RNAi condition. Scale bar: 10 µm. **E**) Representative images of Hela cells mock treated or co-depleted of Zwilch and Bub1, showing that CENP-E is not lost from KTs upon Zwilch and Bub1 co- depletion. Two cells with different depletion efficiency for Zwilch are shown for the RNAi condition. Scale bar: 10 µm.

**Figure 2S3.**
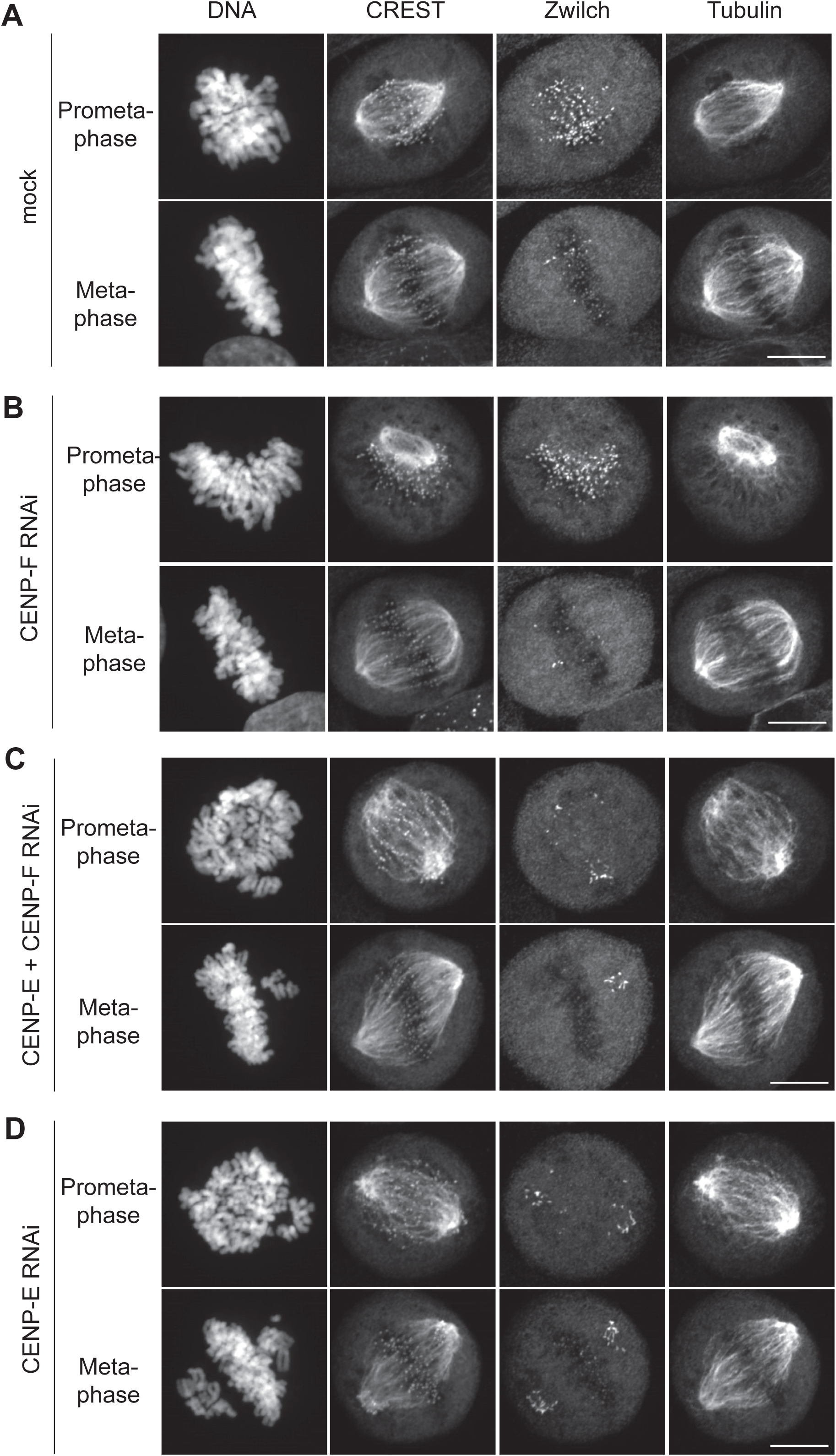
Kinetochore localization of RZZ (Zwilch) is not affected by CENP-E or CENP-F depletion also in presence of microtubules. **A-C**) Representative images of a prometaphase and metaphase HeLa cells either mock treated (A) or individually depleted of CENP-F (B), co-depleted of both CENP-E and CENP-F (C), individually depleted of CENP-E (D) in absence of nocodazole. Neither CENP-E nor CENP-F are required for kinetochore localization of Zwilch even in the presence of spindle microtubules. Scale bar: 10 µm.

**Figure 2S4.**
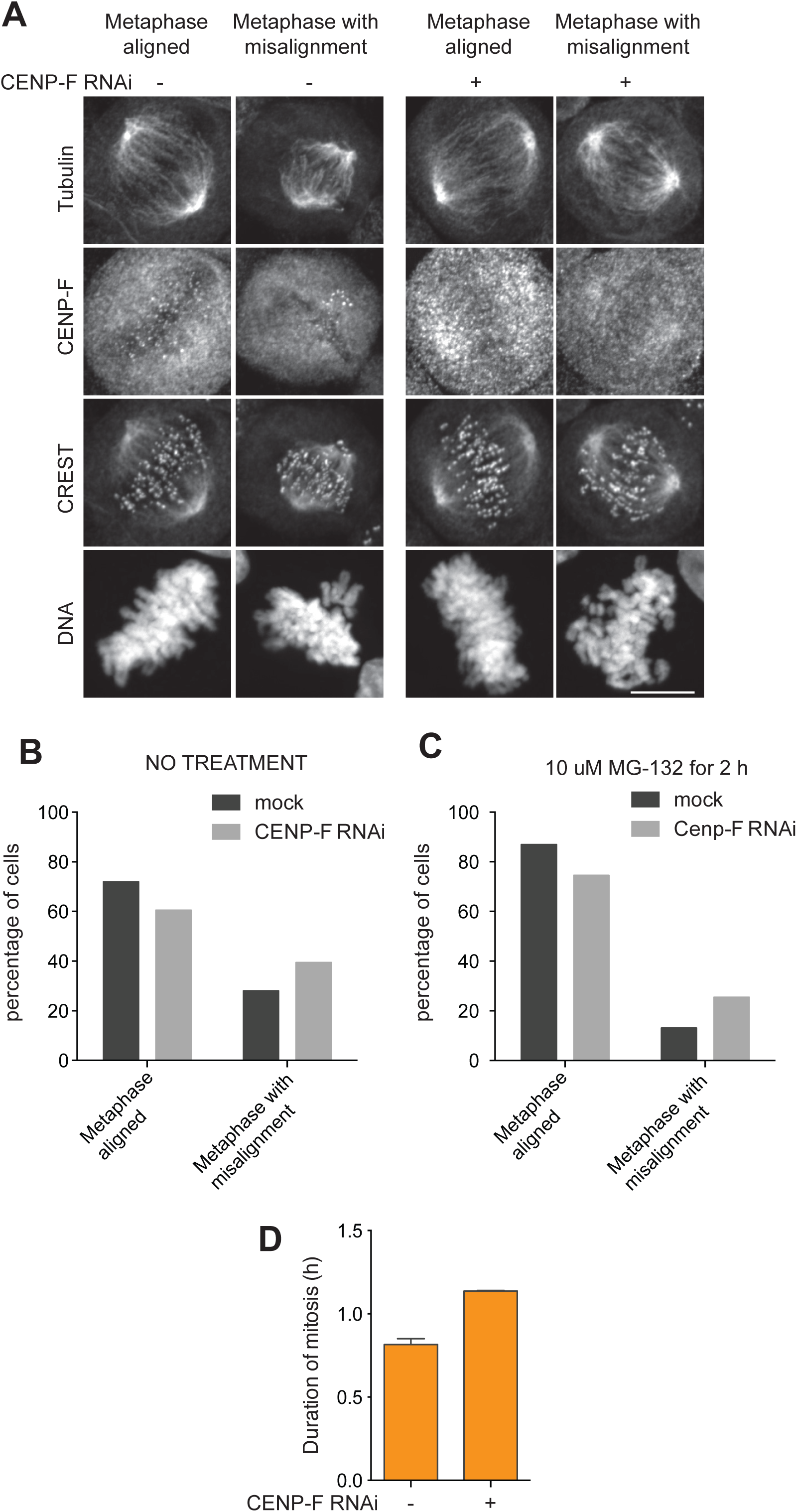
Mitotic phenotype of CENP-F depletion. **A**) Examples of the scored categories, aligned metaphase and metaphases with misalignments in mock treated or CENP-F depleted HeLa cells. Scale bar: 10 µm. **B**) Percentages of cells in the depicted categories in unsynchronized cells from one experiment. **C**) Percentages of cells in the depicted categories in cells treated with 10 µM MG for 2h before fixation from one experiment. **D**) Mean duration of mitosis of HeLa cells in presence or absence of endogenous CENP-F. Cell morphology was used to measure entry into and exit from mitosis by time-lapse microscopy (n>76 per condition per experiment) from two independent experiments. Error bars indicate SEM.

Conversely, depletion of CENP-E or CENP-F in HeLa cells, or even their co-depletion (Figure 2S2A), did not influence the kinetochore localization of Zwilch or Mad1 (Figure 2B-F and Figure 2S3A-D), as observed previously (Martin-Lluesma et al., 2002; Yang et al., 2005). We note, however, that depletion of CENP-E in DLD-1 cells was reported to have deleterious effects on Mad2 localization, while Mad1 or RZZ subunits were not tested (Johnson et al., 2004). Based on these results, we conclude that kinetochore localization of CENP-E and CENP-F does not require the kinetochore corona, nor does it influence corona assembly. We also observed that CENP-E and CENP-F were not reciprocally affected by their depletion (Figure 2G-H), indicating that they localize (at least largely) independently to kinetochores, as previously suggested (Yao et al., 2000).

In most cells analyzed, depletion of CENP-F resulted in apparently normal metaphase alignment, with only a slight increase in the fraction of cells presenting metaphase alignment defects (Figure 2S3B and Figure 2S4). In agreement with the effects of CENP- F depletion being mild, duration of mitosis (caused by spindle assembly checkpoint activation) was only marginally increased in cells depleted of CENP-F (Figure 2S4D). Similarly mild effects from depleting CENP-F were observed previously (Bomont et al., 2005; Feng et al., 2006; Holt et al., 2005; Raaijmakers et al., 2018; Yang et al., 2005). On the other hand, depletion of CENP-E (with or without additional depletion of CENP-F) led to conspicuous chromosome alignment problems (Figure 2S3C-D), as reported previously (Kapoor et al., 2006; Kuhn and Dumont, 2017; Magidson et al., 2011; Magidson et al., 2015; Putkey et al., 2002; Schaar et al., 1997; Wood et al., 1997; Yao et al., 2000; Yen et al., 1991).

### CENP-E binds to the BubR1 pseudokinase domain

Previous studies identified a kinetochore-binding region in residues 2126-2476 of CENP- E (Chan et al., 1998). By expression in insect cells, we generated a recombinant version of a larger fragment of CENP-E (residues 2070-C) encompassing this region fused to eGFP (eGFP-CENP-E^2070-C^) and purified it to homogeneity. After electroporation in mitotic cells arrested by addition of the microtubule-depolymerizing drug nocodazole, cells were fixed to assess the localization of eGFP-CENP-E^2070-C^. eGFP-CENP-E^2070-C^ localized robustly to mitotic kinetochores (Figure 3A), adopting the typical crescent-like shape previously attributed to the corona (Hoffman et al., 2001; Magidson et al., 2015). An equivalent mutant construct in which Cys2697 had been mutated to alanine to prevent farnesylation also localized to kinetochores, even if at generally lower levels and without showing a crescent-like distribution, suggesting that farnesylation is not strictly required for kinetochore localization of CENP-E but that it might contribute to an unknown aspect of corona assembly.

**Figure 3.**
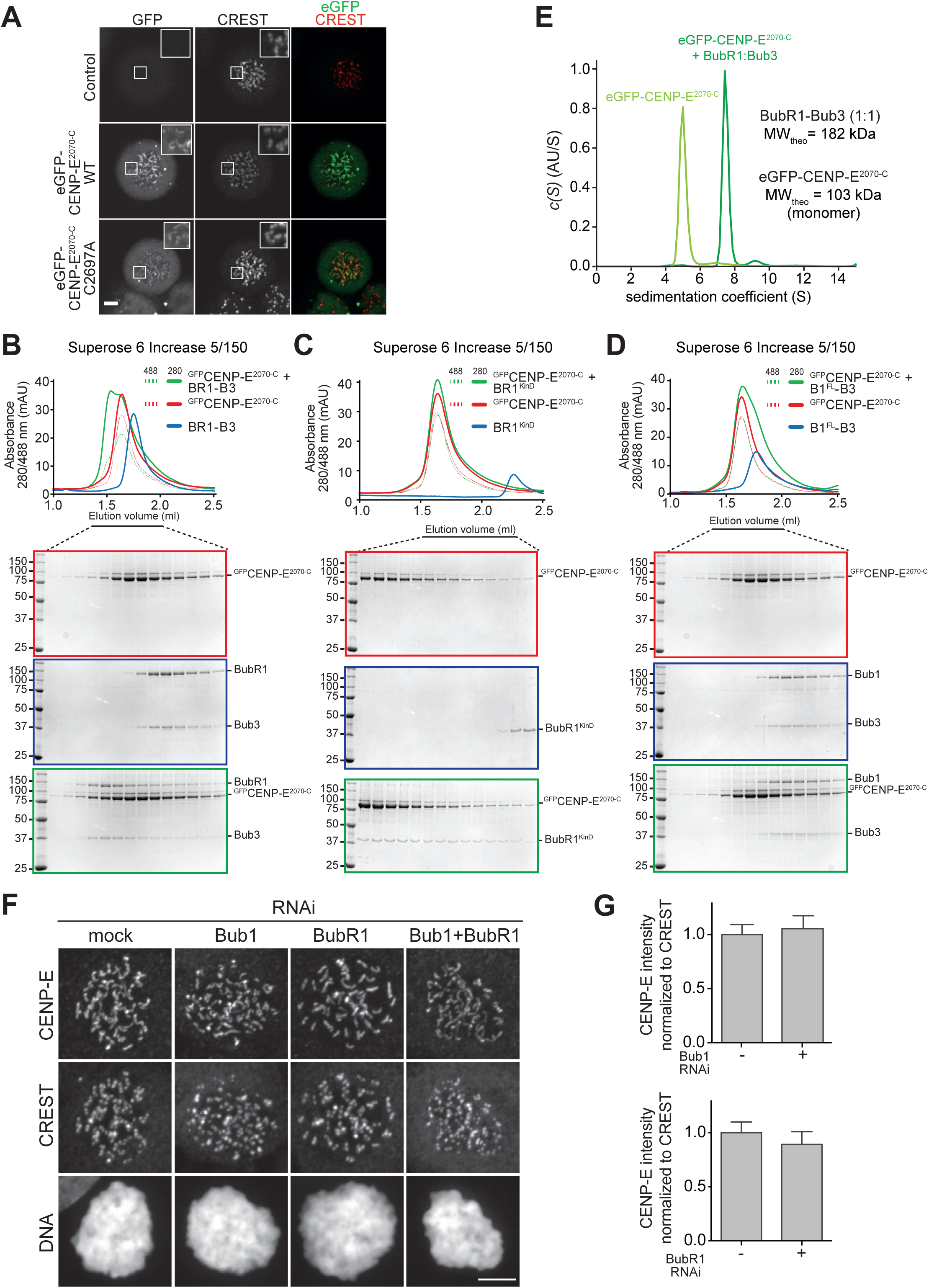
CENP-E interacts with the BubR1 pseudo-kinase domain, but BubR1 is not required for its kinetochore localization. **A**) Representative images of mitotic Hela cells electroporated with eGFP, eGFP-CENP-E^2070-C^ WT or eGFP-CENP-E^2070-C^ C2997A mutant (preventing CENP-E farnesylation). Scale bar: 5 µm. Both the WT and the un-farnesylated mutant CENP-E constructs localize at kinetochores. **B-D**) Elution profiles and SDS-PAGE analysis of SEC experiments of eGFP-CENP-E^2070-C^ with BubR1:Bub3 complex (B), BubR1 pseudo- kinase domain (KinD) construct (C) and Bub1:Bub3 complex (D). A shift in elution volume is observed for the BubR1:Bub3 complex and the BubR1 pseudo-kinase domain construct, indicative of complex formation. The interaction of CENP-E with BubR1 is specific, as no shift is observed with the Bub1:Bub3 complex. **E**) Sedimentation velocity AUC profiles of eGFP-CENP-E^2070-C^ alone and in complex with BubR1:Bub3. AU, arbitrary units; MW_theo_, predicted molecular weight assuming stoichiometry of 1. A reliable estimation of the molecular mass of the proteins in the samples was unsuccessful, likely because of the very elongated and flexible structure of both CENP-E and BubR1. **F**) Representative images of stable Flp-In T-REx cells mock treated or depleted of endogenous Bub1, BubR1, or both, showing that CENP-E kinetochore localization is unaffected under any of the conditions. Scale bar: 10 µm. **G**) Quantification of CENP-E kinetochore levels in cells treated as in (F). The graph shows mean intensity of two independent experiments, the error bars indicate SEM. The mean value for non-depleted cells expressing GFP was set to 1.

**Figure 3S1.**
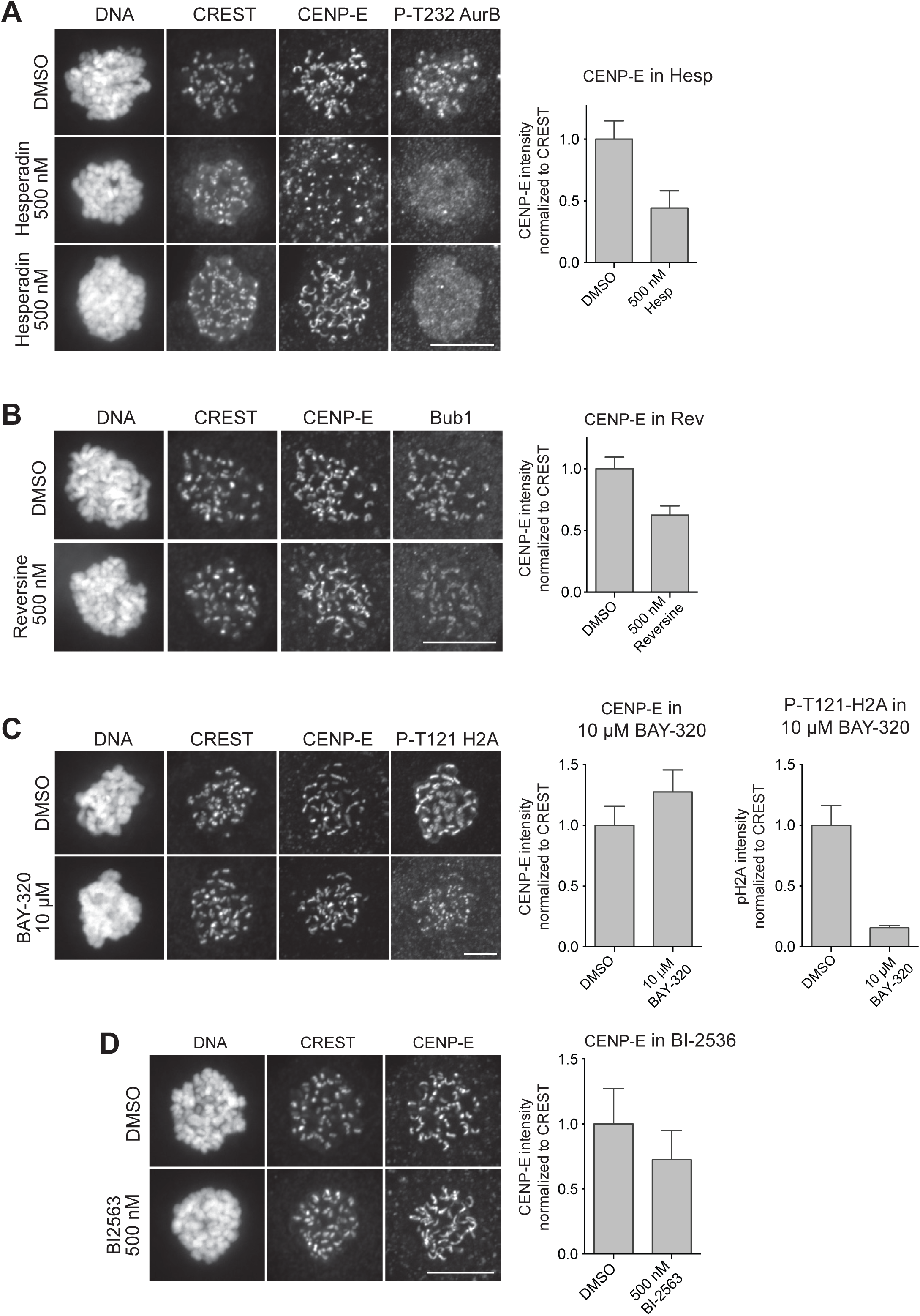
CENP-E kinetochore localization sensitivity to kinases inhibition. **A-D**) Representative images and quantification of CENP-E kinetochore levels in mitotic Hela cells treated with the indicated concentrations of the indicated kinase inhibitors. Scale bar: 10 µm. CENP-E localization is sensitive to Aurora B (A), Mps1 (B) and, to a lesser extent, Plk1 (D) but not to Bub1 (C) inhibition. Reduction in P-T232 Aurora B (Aurora B activation segment) staining was used as a positive control for Aurora B inhibition (A), while reduction in Bub1 localization was used as positive control for Mps1 inhibition (B). Bub1 inhibition was confirmed by reduction in P-T121 H2A staining (C). The graphs show mean intensity of one (C, I), two (A) or four (B) experiments. The error bars indicate SEM and the mean values for DMSO-treated cells are set to 1.

In previous yeast 2-hybrid (Y2H) analyses, a CENP-E segment encompassing residues 1958-2662 was found to interact with residues 410-1050 of BubR1 (Chan et al., 1998; Yao et al., 2000). Even if shorter by more than 100 residues at the N-terminal end, eGFP-CENP-E^2070-C^ interacted directly with the dimeric BubR1:Bub3 complex in size exclusion chromatography (SEC) analyses (Figure 3B), as evidenced by the shift in elution volume of both proteins when combined at 16 µM and 4 µM concentration, respectively. Similar observations were made when we mixed CENP-E^2070-C^ with the BubR1 pseudokinase domain (KinD, residues 705–1050) (Figure 3C). eGFP-CENP- E^2070-C^, however, did not interact with the paralogous Bub1:Bub3 dimer (Figure 3D). In analytical centrifugation (AUC) sedimentation velocity experiments in which we monitored the sedimentation of eGFP-CENP-E, addition of unlabeled BubR1:Bub3 at 3-fold higher concentration caused a complete shift of eGFP-CENP-E to a species with higher sedimentation coefficient (S), indicative of complex formation (Figure 3E). The high frictional ratio of this sample (an indication that the CENP-E structure is very elongated, a consequence of its large coiled-coil content) prevented a quantitative estimate of molecular mass. The analysis, however, strongly suggest that eGFP-CENP- E^2070-C^ adopts the highly elongated conformation of coiled-coils, as shown previously for recombinant full length CENP-E from *Xenopus laevis* (Kim et al., 2008). Thus, a minimal segment of CENP-E capable of kinetochore localization interacts directly with the BubR1:Bub3 complex, and the BubR1 pseudokinase domain is sufficient for this interaction, at least at the relatively high concentration of the SEC assay. In agreement with our own previous studies (Breit et al., 2015), BubR1 did not show any catalytic activity, nor did it become active in presence of eGFP-CENP-E^2070-C^ (unpublished data).

In a previous study in egg extracts of *Xenopus laevis*, depletion of BubR1 was shown to prevent kinetochore localization of CENP-E, an effect that could be rescued by re-addition of wild type BubR1, but not of a deletion mutant lacking the kinase domain (Mao et al., 2003). CENP-E kinetochore levels were also reduced in DLD-1 cells upon depletion of BubR1 by RNAi (Johnson et al., 2004).

We therefore asked if BubR1 was also important for CENP-E recruitment in HeLa cells. Furthermore, in view of evidence that Bub1 is required for kinetochore recruitment of BubR1 (Klebig et al., 2009; Liu et al., 2006; Logarinho et al., 2008; Overlack et al., 2015), we also monitored localization of CENP-E upon depletion of Bub1. Contrarily to the previous observations in frogs and DLD-1 cells, but in agreement with other studies in HeLa cells (Akera et al., 2015; Lampson and Kapoor, 2005; Liu et al., 2006), RNAi-based depletion of Bub1 or BubR1 did not result in obvious adverse effects on the kinetochore localization of CENP-E, even after co-depletion of Zwilch (Figure 3F-G and Figure 2S2B-E). These observations suggest that CENP-E, at least in HeLa cells, becomes recruited through a different pathway that does not involve Bub1 and BubR1. After application of highly specific small-molecule inhibitors, we found CENP-E kinetochore localization to depend on the kinase activity of Aurora B and (to a lower extent) of Mps1, but not of Bub1 or of Plk1 (Figure 3S1A-D). The dependence of CENP-E on Aurora B kinase activity for kinetochore localization, together with the central spindle co-localization at anaphase of CENP-E with the chromosome passenger complex (CPC, the catalytic subunit of which is Aurora B), leads to speculate that these proteins interact, a hypothesis that will need to be formally tested in the future.

### CENP-F binds to the Bub1 kinase domain

Next, we asked how CENP-F becomes recruited to kinetochores. CENP-F recruitment was strictly dependent on the kinase activity of Aurora B, partly dependent on that of Mps1 and Plk1, and not dependent on that of Bub1 (Figure 4A and Figures 4S1). This pattern of kinetochore localization is reminiscent of that of Bub1, which has been previously shown to be important for CENP-F kinetochore recruitment (Johnson et al., 2004; Klebig et al., 2009; Liu et al., 2006; Raaijmakers et al., 2018). In agreement with these previous studies, RNAi-based depletion of Bub1 resulted in complete ablation of CENP-F from kinetochores (Figure 4C, see panel I for quantification).

**Figure 4.**
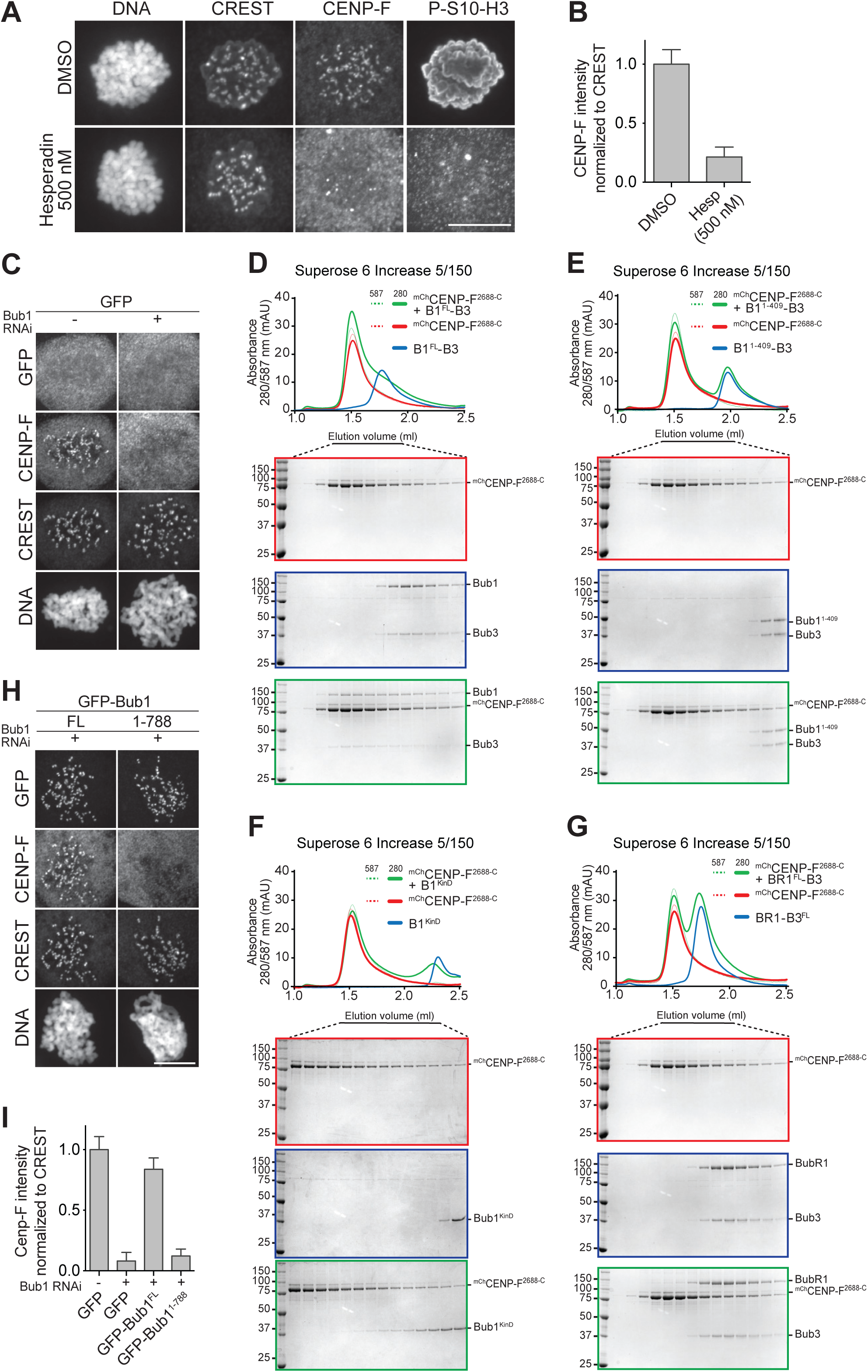
CENP-F interaction with the Bub1 kinase domain is necessary for its kinetochore localization. **A**) Representative images of mitotic HeLa cells treated with 500 nM Hesperadin, showing that CENP-F kinetochore localization is strictly dependent on Aurora B kinase activity. Reduction in P-S10-H3 staining was used as a control for the Aurora B inhibition. Scale bar: 10 µm. **B**) Quantification of CENP-F kinetochore levels in cells treated as in A. The graph shows mean intensity of three experiments. The error bars indicate SEM and the mean values for DMSO-treated cells are set to 1. **C**) Representative images of GFP-expressing stable HeLa Flp-In T-REx cell lines mock treated or depleted of Bub1, showing that CENP-F kinetochore recruitment depends on the presence of Bub1 at kinetochores. **D-G**) Elution profiles and SDS-PAGE analysis of SEC experiments of mCherry-CENP-F^2688-C^ with the Bub1^FL^:Bub3 complex; FL, full length (D), the Bub1^1-409^:Bub3 complex (E), the Bub1 kinase domain (KinD) (F), and the BubR1:Bub3 complex (G). A shift in the elution volume is only observed for the Bub1 constructs containing the C-terminal kinase domain (D, F) The interaction of CENP-F with Bub1 is specific, as no shift is observed for the BubR1:Bub3 complex (G). **H**) Representative images of stable HeLa Flp-In T-REx cell lines depleted of endogenous Bub1 and expressing GFP-Bub1 full length or lacking the kinase domain (Bub1^1-788^). CENP-F kinetochore recruitment depends on the Bub1 kinase domain, as Bub1^1-788^ does not rescue CENP-F localization, while full length Bub1 does. Scale bar: 10 µm. **I**) Quantification of CENP-F kinetochore levels in cells of panels C and H. The graph shows mean intensity of three independent experiments, the error bars indicate SEM. The mean value for non-depleted cells expressing GFP is set to 1.

**Figure 4S1.**
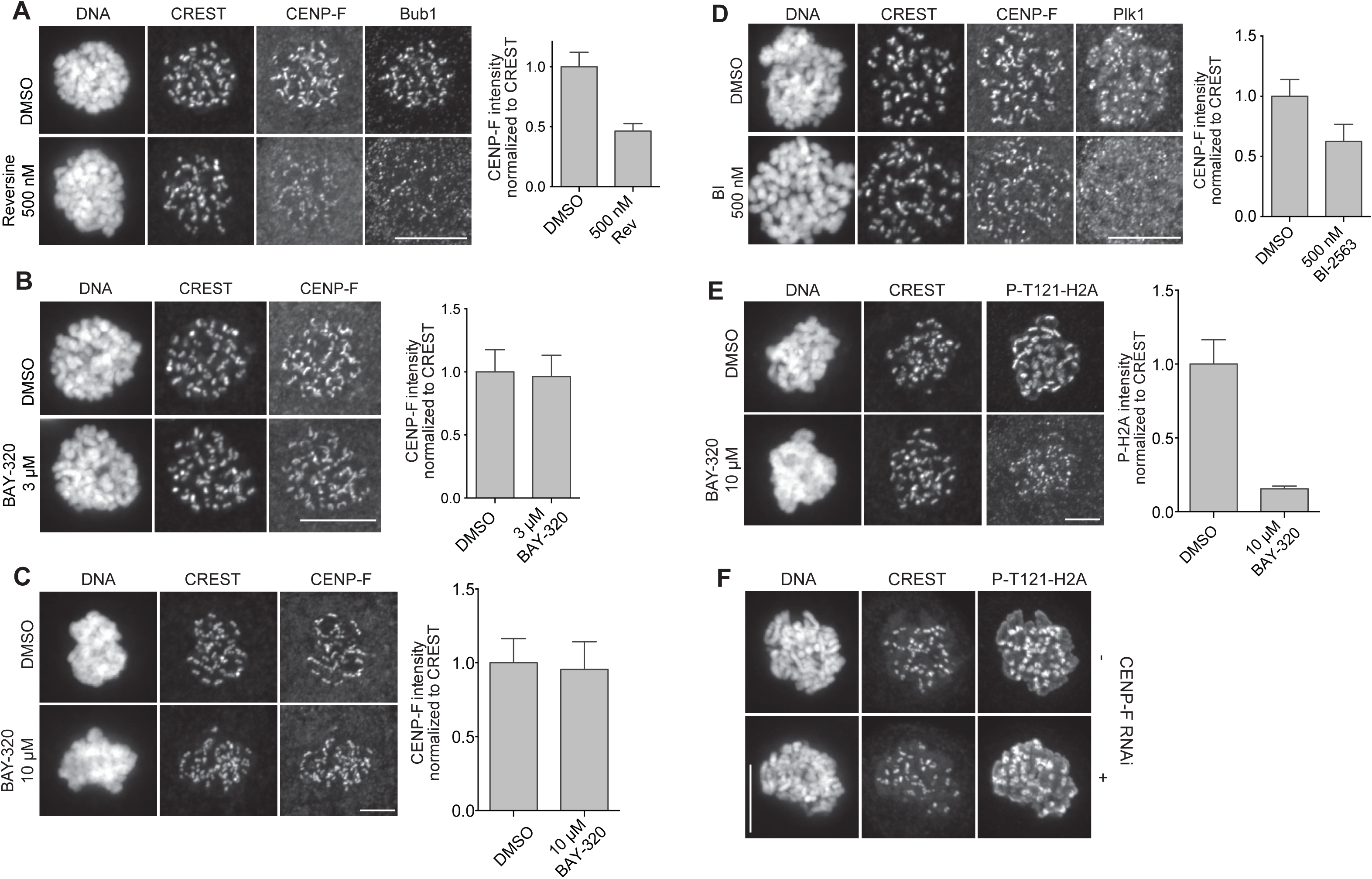
Sensitivity to kinases inhibition of CENP-F kinetochore localization. **A-E**) Representative images and quantification of CENP-F kinetochore levels in mitotic Hela cells treated with the indicated type and concentrations of kinase inhibitors. Scale bar: 10 µm. CENP-F localization is sensitive to Mps1 (A) and partially to Plk1 (D) inhibition. Despite a requirement for the Bub1 kinase domain for CENP-F kinetochore recruitment, Bub1 catalytic activity is dispensable (B, C). Reduction in Bub1 localization was used as a control for Mps1 inhibition (A) and reduction in P-T121-H2A staining was used as a control for Bub1 inhibition (E). Loss of Plk1 localization was used as control for Plk1 inhibition (D). The graphs show mean intensity of one (B, C, E) or two (A, D) experiments. The error bars indicate SEM and the mean values for DMSO-treated cells are set to 1. **F**) Representative images of HeLa cells mock treated or depleted of CENP-F showing that CENP-F depletion does not affect the kinase activity of Bub1, as no changes are detected for the P-T121-H2A signal. Scale bar: 10 µm.

By expression in insect cells, we generated a recombinant fragment of CENP-F encompassing its previously identified kinetochore-binding domain (residues 2688-C) (Hussein and Taylor, 2002; Zhu, 1999; Zhu et al., 1995a) fused to an N-terminal mCherry tag. In SEC experiments, mCherry-CENP-F^2688-C^ bound Bub1:Bub3 directly, as indicated by its altered elution volume in presence of the CENP-F construct (Figure 4D; note that mCherry-CENP-F^2688-C^ is highly elongated, as shown below, and therefore its hydrodynamic radius, which determines elution volume in SEC experiments, is unlikely to change as a result of an interaction with Bub1:Bub3). On the other hand, mCherry-CENP-F^2688-C^ failed to interact with Bub1^1-409^:Bub3, where the Bub1 deletion mutant Bub1^1-409^ lacks a central region of Bub1 and its kinase domain (Figure 4E). Indeed, mCherry-CENP-F^2688-C^ bound the Bub1 kinase domain (Bub1^KinD^, residues 725–1085; Figure 4F). Conversely, mCherry-CENP-F^2688-C^ did not interact with the BubR1:Bub3 complex (Figure 4G). Thus, the kinetochore-targeting domain of CENP-F interacts directly with the Bub1:Bub3 complex, and the kinase domain appears to be necessary and partly sufficient for this interaction.

In agreement with these *in vitro* findings, we observed robust kinetochore localization of endogenous CENP-F in HeLa cells previously depleted of Bub1 by RNAi and expressing an RNAi-resistant GFP-Bub1 transgene, while CENP-F kinetochore localization appeared entirely compromised in Bub1-depleted cells expressing GFP-Bub1^1-788^, which lacks exclusively the Bub1 kinase domain (Figure 4H-I). Collectively, our observations indicate that the kinase domain of Bub1 is sufficient for a direct interaction with CENP- F *in vitro*, and necessary for kinetochore recruitment of CENP-F in HeLa cells. A very recent study identified a similar requirement for the kinase domain of Bub1 in kinetochore recruitment of CENP-F in HAP1 cells (Raaijmakers et al., 2018).

### CENP-F dimerization is important for Bub1 binding

The mCherry-CENP-F^2688-C^ construct that interacted with Bub1:Bub3 in SEC experiments also localized to mitotic kinetochores when electroporated in HeLa cells (Figure 5A). A farnesylation mutant of this construct on which Cys3207 had been mutated to alanine retained kinetochore localization, although not as robustly as the wild type counterpart (Figure 5A). This result suggests that farnesylation is not strictly required for kinetochore recruitment of CENP-F, as already observed with CENP-E (Figure 3A). Collectively, our results with electroporated farnesylation mutants of CENP- E and CENP-F are in agreement with results obtained with farnesyl transferase inhibitors, in which only partial repression of kinetochore recruitment of CENP-E and CENP-F was observed (Holland et al., 2015).

**Figure 5.**
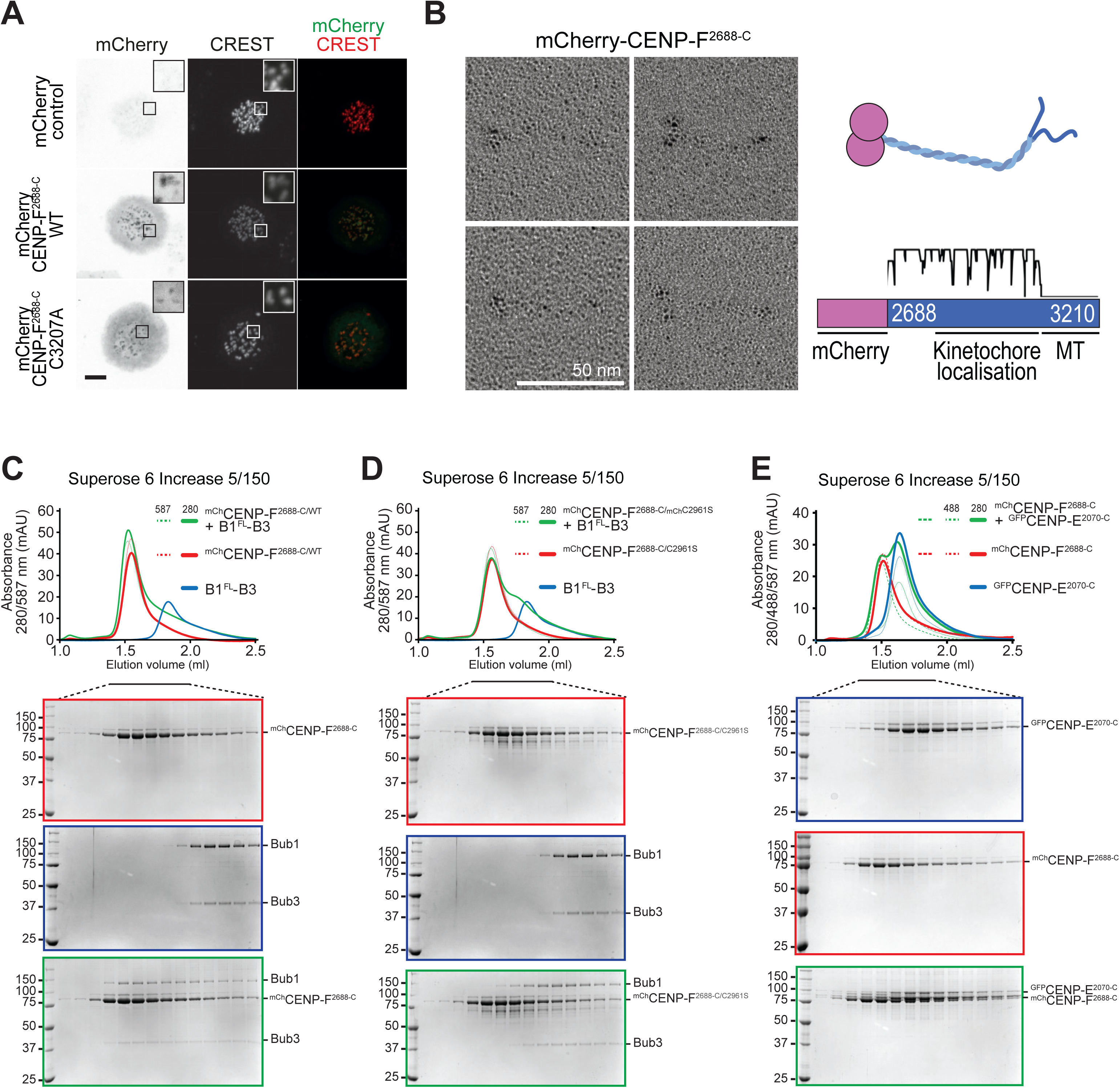
Requirements for CENP-F kinetochore localization. **A**) Representative images of mitotic HeLa cells electroporated with mCherry, mCherry-CENP-F^2688-C^ WT or mCherry-CENP-F^2688-C^ C3207A (farnesylation) mutant. Scale bar: 5µm. As for CENP-E, both the WT and the un-farnesylated mutant CENP-F constructs localize at kinetochore. **B**) mCherry-CENP-F^2688-C^ sample was visualized by electron microscopy after glycerol spraying and low-angle platinum shadowing (right panel). The elongated shape of the observed particles is consistent with the secondary structure expected for the mCherry-tag coiled-coil construct (right panel). **C-D**) SEC elution profiles and SDS-PAGE analysis of binding experiments with 16 µM each of mCherry-CENP-F^2688-C^ WT (C) or C2961S mutant (D) and 4 µM Bub1^FL^:Bub3 complex. The shift in elution volume of Bub1:Bub3 is observed with both wild type and mutant CENP-F, but it significantly less pronounced for the CENP-F mutant, suggesting that the C2961S mutation reduces the affinity of CENP-F for the Bub1 kinase domain without completely abolishing it. **E**) SEC elution profile and SDS-PAGE analysis of a binding experiment with 16 µM each of mCherry-CENP-F^2688-C^ and eGFP-CENP-E^2070-C^. No shift is observed, indicating that the tested constructs do not interact.

By rotary shadowing electron microscopy (EM), which is particularly suited to the study of elongated coiled-coil proteins, mCherry-CENP-F^2688-C^ had the appearance of a highly elongated (∼40 nm) rod. Most likely, the latter corresponds to a predicted coiled-coil comprised between residues 2688 and ∼3000, flanked on one side by two globular domains, most likely corresponding to mCherry, and on the other side by disordered fragments corresponding to the last ∼200 residues and containing the C-terminal microtubule-binding domains (Figure 5B) (Feng et al., 2006; Musinipally et al., 2013; Volkov et al., 2015). Collectively, these observations suggest that CENP-F^2688-C^ contains a parallel dimeric coiled-coil, like the one previously identified in CENP-E (Kim et al., 2008).

On the basis of previous studies implicating Cys^2864^ in kinetochore recruitment of mouse CENP-F (Zhu, 1999), we generated a mutant version of human mCherry-CENP-F^2688-C^ in which the equivalent residue, Cys^2961^, was mutated to serine (our residue numbering is in accordance with the 3210-residue human CENP-F sequence in Uniprot). While wild type mCherry-CENP-F^2688-C^ interacted with Bub1:Bub3 in SEC experiments, as already shown, the interaction was at least partially impaired when the mCherry-CENP-F^2688^- C/C^2961S^ mutant was analyzed, confirming the role of Cys2961 in kinetochore localization and implicating this residue in the interaction with Bub1 (Figure 5C-D). Furthermore, mCherry-CENP-F^2688-C^ did not interact with GFP-CENP-E^2070-C^ in SEC experiments, as predicted based on previous work identifying the CENP-E-binding region of CENP-F within a segment (residues 1804-2104) that precedes and is not included in CENP-F^2688-C^ (Figure 5E) (Chan et al., 1998; Yao et al., 2000).

The effects of the Cys^2961^ mutation made us ask if we could identify a minimal Bub1-binding domain of CENP-F. For this, we further trimmed CENP-F. CENP-F^2866-2990^, which is entirely encompassed within the predicted coiled-coil of CENP-F, appeared dimeric by sedimentation velocity AUC and retained the ability to bind to the Bub1 kinase domain in a SEC experiment (Figure 6A-B). Similar results were obtained with an even shorter CENP-F fragment, CENP-F^2922-2990^ (Figure 6C-D). CENP-F^2950-2990^, on the other hand, appeared monomeric in AUC runs and was unable to interact with the Bub1 kinase domain (Figure 6E-F). These observations do not allow us to resolve whether impaired binding to the Bub1 kinase domain CENP-F^2950-2990^ is due to loss of dimerization or to trimming of residues directly involved in the interaction, but identify CENP-F^2922-2990^ as a minimal Bub1-binding fragment of CENP-F. In agreement with these observations, mCherry-CENP-F^2866-2990^ and mCherry-CENP-F^2922-2990^ labeled kinetochores after electroporation in HeLa cells, albeit weakly in comparison to mCherry-CENP-F^2866-C^, whereas mCherry-CENP-F^2950-2990^ did not localize to kinetochores (Figure 6G).

**Figure 6.**
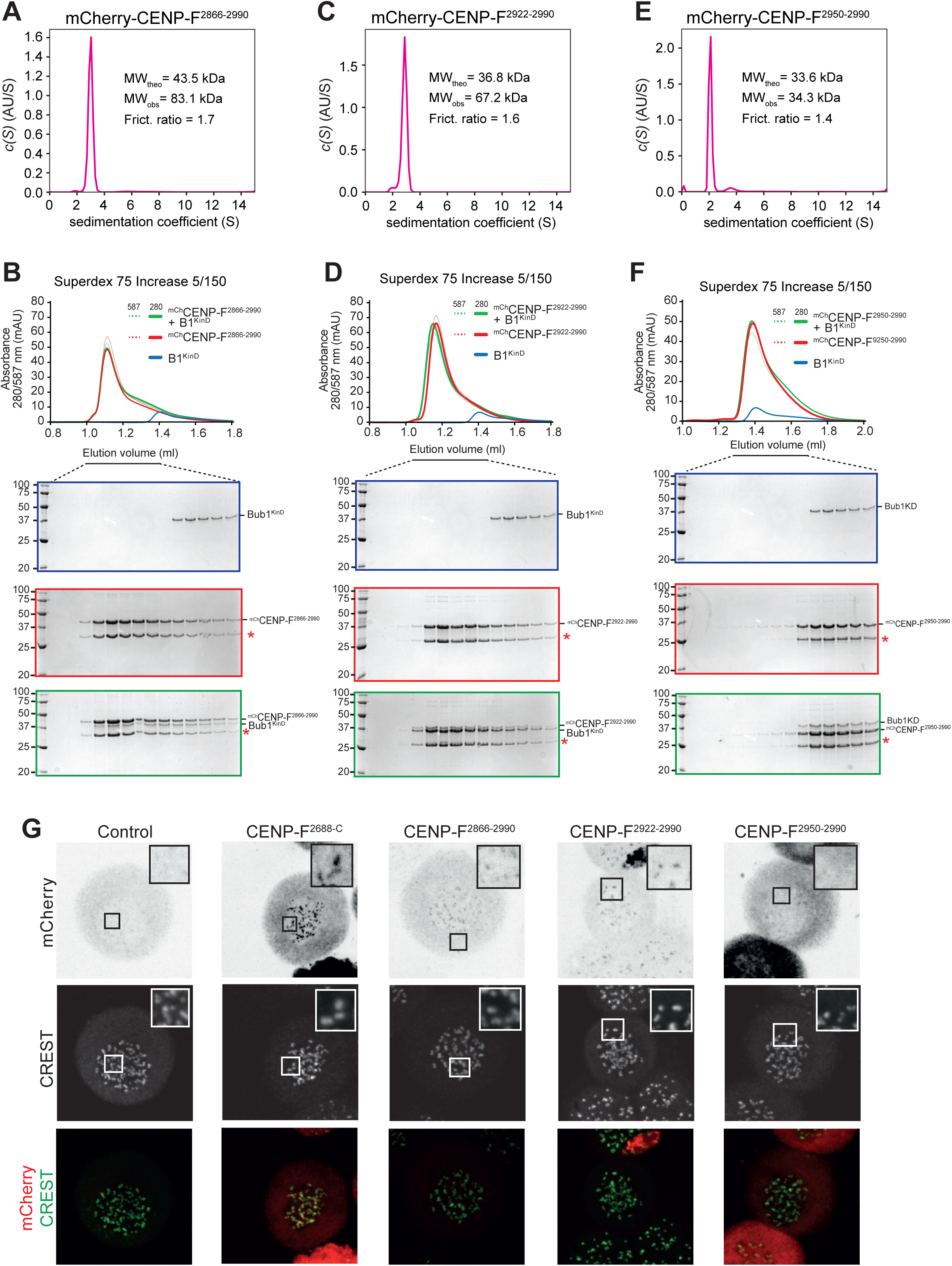
Identification of a minimal CENP-F construct for binding to Bub1. **A**) Sedimentation velocity AUC results of the indicated mCherry-CENP-F constructs. MW_obs_, observed molecular weight; MW_theo_, the predicted molecular weight of the monomer; Frict. ratio denotes the frictional ratio. AU, arbitrary units. mCherry-CENP- F^2866-2990^ forms a dimer. **B**) Elution profiles and SDS-PAGE analysis of SEC experiments of the Bub1 kinase domain (KinD) with mCherry-CENP-F^2866-2990^. The shift in elution volume of Bub1^KinD^ indicated binding. The red asterisk indicated a breakdown product of mCherry that is produced during boiling in sample buffer. **C**) As in (A) but with the CENP-F^2922-2990^ construct, which is also dimeric. **D**) As in (B) but with the CENP-F^2922-2990^ construct. Also in this case, an interaction with the kinase domain of CENP-F is clearly discernible. **E**) As in (A) but with the CENP-F^2950-2990^ construct, which is monomeric. **F**) As in (B) but with the CENP-F^2950-2990^ construct. In this case, no interaction with the kinase domain of CENP-F is discernible. **G**) Representative images of mitotic HeLa cells electroporated with the indicated constructs. mCherry-CENP-F^2688-C^ (positive control), mCherry-CENP-F^2866-2990^, and mCherry-CENP-F^2922-2990^ localized to kinetochores, whereas mCherry (negative control) and mCherry-CENP-F^2950-2990^ did not. Scale bar: 5µm.

## Conclusions

Previous studies on human Bub1 and BubR1, including our own work, demonstrated that these paralogs sub-functionalized in various ways, including 1) the selective inactivation of the kinase domain in BubR1; 2) the development of phospho-aminoacid recognition modules that contribute to the ability of Bub3 to recognize distinct substrates; and 3) the interaction with distinct binding partners that subtends to distinct functions in chromosome alignment and mitotic checkpoint signaling (Figure 7) (Overlack et al., 2017; Overlack et al., 2015; Primorac et al., 2013; Suijkerbuijk et al., 2012; van Hooff et al., 2017).

**Figure 7.**
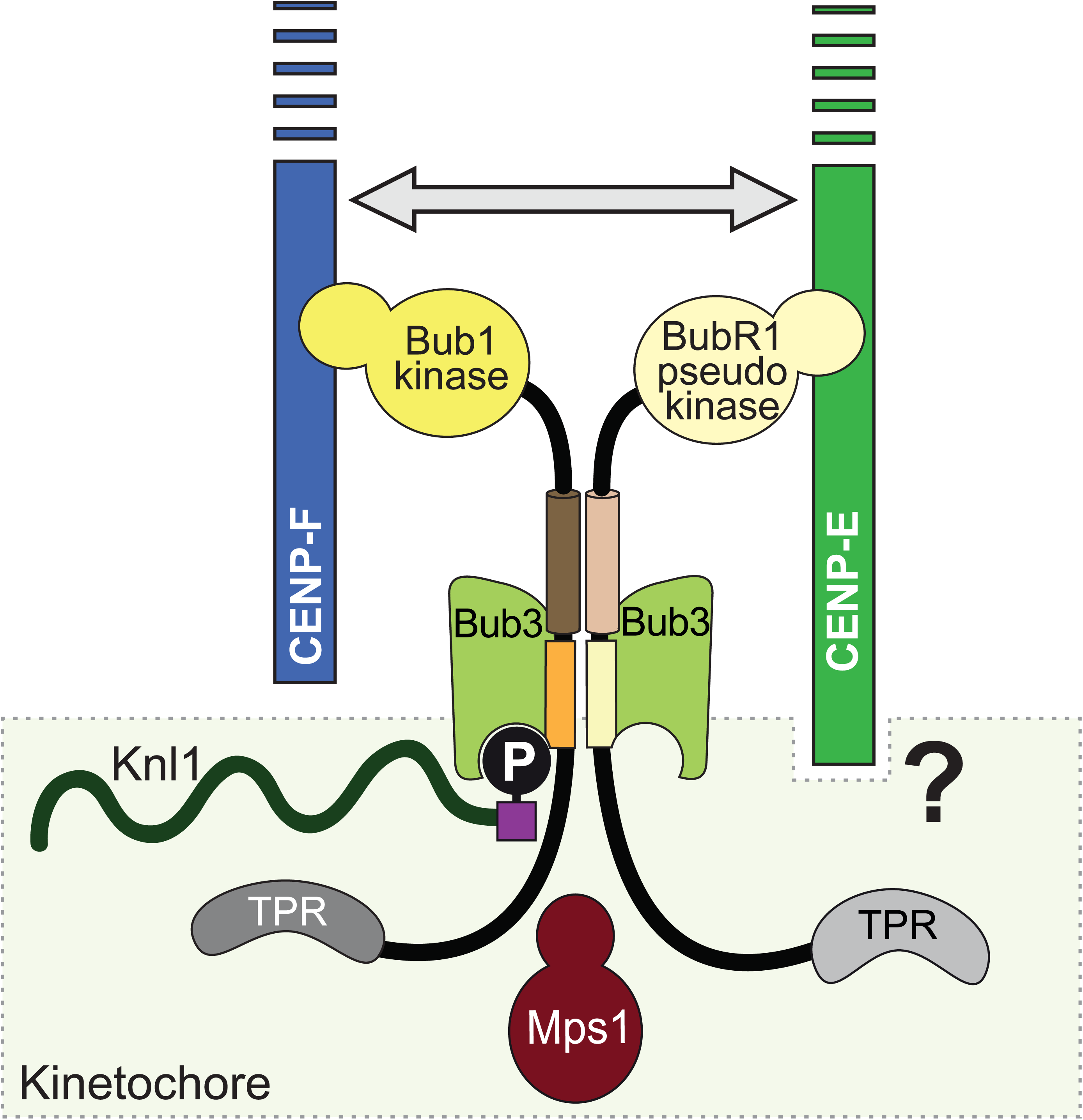
Schematic of the interactions of Bub1 and BubR1 paralogs. The schematic summarizes the interactions occurring at kinetochores between CENP-E, CENP-F, Bub1, and BubR1. Bub1 is recruited to the kinetochore subunit Knl1 (see Introduction) after phosphorylation by the SAC kinase Mps1. There, Bub1 recruits BubR1 through a pseudo-dimeric interaction (Musacchio, 2015). CENP-F kinetochore localization strictly depends on Bub1, while CENP-E recruitment requires a wider and still uncharacterized network of interactions, indicated by a question mark. RZZ and Mad1 recruitment, as well as the corona expansion, appear to be independent from CENP-E and CENP-F, and are not shown. An interaction of CENP-E and CENP-F has also been identified (grey arrow), but is not sufficient for CENP-F localization in absence of Bub1.

In this study, we report an additional aspect of this sub-functionalization, and show that the kinase domains of BubR1 and Bub1 interact respectively with C-terminal regions of CENP-E and CENP-F that encompass the kinetochore-targeting domains of these proteins. Both interactions are direct and were reproduced with recombinant proteins. Neither interaction appears to be crucially required for downstream signaling events. In the BubR1 case, it appears well established that deletion of the pseudokinase domain in human cells is compatible with its functions in the SAC as a subunit of the MCC, the SAC effector (Musacchio, 2015). We, and others, have also shown that depletion of BubR1 does not affect CENP-E kinetochore localization in HeLa cells [this study and (Akera et al., 2015; Lampson and Kapoor, 2005)]. In *Xenopus* egg extracts, however, the kinase domain of BubR1 has been implicated in CENP-E recruitment and this interaction has been shown to be important for SAC silencing (Mao et al., 2003). Given the very complex evolutionary history of the Bub1 and BubR1 paralogs or of the singleton from which they originate, these apparent differences may genuinely reflect different evolutionary paths in these organisms (Suijkerbuijk et al., 2012; van Hooff et al., 2017). A question for future work that this study raises regards the detailed mechanism of kinetochore recruitment of CENP-E in human cells, which remains unknown.

The role of Bub1 in CENP-F kinetochore recruitment was already established in previous work (Berto et al., 2018; Johnson et al., 2004; Liu et al., 2006; Raaijmakers et al., 2018), and a very recent study implicated the Bub1 kinase domain in CENP-F kinetochore recruitment (Raaijmakers et al., 2018). Here, we have extended this previous study by showing that the interaction of the Bub1 kinase domain and CENP-F is direct, and by identifying a minimal CENP-F domain involved in this interaction and capable of kinetochore localization. Our identification of a minimal kinetochore-targeting domain of HsCENP-F within residues 2922-2990 agrees with a previous study that made use of CENP-F deletion mutants (Zhu, 1999). It also agrees with another study, currently in press, that identified distinct binding domains in CENP-F for nuclear envelope and for kinetochore localization (Berto et al., 2018). Specifically, residues 2655 to 2860 of mouse CENP-F (corresponding to residues 2866-3072 of HsCENP-F) were sufficient for kinetochore recruitment (Berto et al., 2018). Within this fragment, a specialized N-terminal sub-domain (residues 2655-2723, corresponding to HsCENP-F residues 2866-2933) bound Nup133, a subunit of the nuclear pore complex Nup107-Nup160, and mediated CENP-F recruitment to the nuclear envelope shortly before mitosis, but was not required for kinetochore recruitment. A specialized C-terminal sub-domain (residues 2724-2860, corresponding to HsCENP-F residues 2934-3072), on the other hand, was required for kinetochore recruitment (Berto et al., 2018). As previous studies identified CENP-E as binding partner of CENP-F (Chan et al., 1998; Mao et al., 2003; Yao et al., 2000), CENP-E may reinforce binding of CENP-F to kinetochores, as shown by Berto and colleagues (Berto et al., 2018). However, there is now sufficient evidence to conclude that this interaction is clearly not sufficient to promote stable binding of CENP-F to kinetochores in absence of Bub1.

Our observation that CENP-F depletion results in very mild chromosome alignment defects, in line with other reports (see e.g. (Raaijmakers et al., 2018)), is surprising. CENP-F has been implicated in Dynein recruitment and regulation through a pathway involving Nde1, Ndel1, and Lis1, the product of the lissencephaly type 1 gene (Baffet et al., 2015; Bolhy et al., 2011; Hu et al., 2013; Simoes et al., 2018; Vergnolle and Taylor, 2007). However, CENP-F is not sufficient for stable kinetochore recruitment of Dynein, as it does not seem to be able to complement the very strong reduction or loss of kinetochore Dynein in cells depleted of the RZZ complex or Spindly [see for instance (Barisic et al., 2010; Chan et al., 2009; Gassmann et al., 2010)]. The latter appears therefore to be the dominant factor in Dynein recruitment to kinetochores. It is plausible, however, that the consequences of CENP-F depletion are exacerbated by concomitant depletion of RZZ (Simoes et al., 2018).

As already discussed in the Introduction, various common features of CENP-E and CENP-F support the speculation that they are distantly related paralogs. Their ability to interact with the kinase domains of BubR1 and Bub1, themselves paralogs, lends strong further credit to this hypothesis. Apparent lack of strong functional consequences from disrupting these interactions may indicate that they may have become vestigial in some species. It is also possible, however, that these interactions play more important roles during development or in specific cell types. The dissection described here will allow testing this hypothesis in future work.

## Acknowledgments

We thank all members of the Musacchio laboratory for helpful discussions and comments. A.M. gratefully acknowledges the Max Planck Society, the European Research Council (ERC) Advanced Investigator Grant RECEPIANCE (proposal number n° 669686), and the European Union’s Horizon 2020 research and innovation programme under the Marie Sklodowska-Curie grant agreement No 675737. We are very grateful to Dr. Valerie Doye and her collaborators for sharing with us unpublished results.

## Material & Methods

### Plasmids

The codon optimized cDNAs of *Homo sapiens* CENP-E (Q02224) and CENP- F (P49454) were synthetized at GeneWiz. CENP-E and CENP-F constructs were subcloned respectively in pLIB-eGFP and pLIB-mCherry or pET-mCherry, modified versions of the pLIB (Weissmann et al., 2016) and pET-28 vectors for expression of proteins with N-terminal PreScission-cleavable His_6_-eGFP or His_6_-mCherry tag. Site-directed mutagenesis was performed by PCR (Sawano and Miyawaki, 2000). All constructs were sequence verified. The vectors for the co-expression of full length Bub1 and BubR1 proteins with Bub3, as well as that for the Bub1 & BubR1 constructs were described previously (Breit et al., 2015; Overlack et al., 2015).

### Protein expression and purification

Expression and purification of eGFP-CENP- E^2070-C^ and mCherry-CENP-F^2866-C^ wild-type and mutants was carried out in insect cells using a pBig system (Weissmann et al., 2016). Baculoviruses were generated in Sf9 cells and use to infect Tnao38 cells for 48-96 hours at 27°C. Cells were collected by centrifugation, washed in PBS and then frozen at −80°C. CENP-E expressing cell pellets were resuspended in lysis buffer (50 mM Sodium Phosphate buffer pH 8.0, 500 mM NaCl, 5 % (w/v) glycerol and 0.5 mM TCEP) supplemented with protease inhibitor cocktail, lysed by sonication and cleared by centrifugation at 100.000 g at 4°C. The supernatant was filtered and loaded on a 5 ml HisTrap FF column (GE Healthcare) equilibrated in lysis buffer. After washing with lysis buffer, the protein was eluted with a linear gradient of 0-250 mM imidazole in 10 column volumes. The fractions of interest were pooled, concentrated with a 50 kDa cut-off Amicon concentrator (Millipore) and loaded onto a Superose 6 Increase 10/300 (GE Healthcare) equilibrated in SEC buffer (50 mM Hepes pH 8.0, 200 mM NaCl, 5% (w/v) glycerol and 0.5 mM TCEP). CENP-E containing fractions were concentrated, flash-frozen in liquid nitrogen and stored at - 80°C. The purification protocol for the mCherry-CENP-F^2866-C^ constructs is identical to that of eGFP-CENP-E^2070-C^, but the lysis and the SEC buffers were at pH 7.0.

The constructs mCherry-CENP-F^2866-2990^, mCherry-CENP-F^2922-2990^ and mCherry-CENP- F^2950-2990^ were expressed in *E. coli* BL21 (DE3) RP plus cells grown at 37°C to O.D._600_ = 2 and then induced with 0.25 mM IPTG for 16 h at 25°C. Cell were collected by centrifugation, washed in PBS and then frozen at −80°C. Cell pellets were resuspended in lysis buffer (50 mM Sodium Phosphate buffer pH 7.5, 500 mM NaCl, 5 % (w/v) glycerol and 2 mM β-mercaptoethanol) supplemented with protease inhibitor cocktail, lysed by sonication and cleared by centrifugation at 70.000 g at 4°C. The supernatant was filtered and loaded on a 5 ml HisTrap FF column (GE Healthcare) equilibrated in lysis buffer. After washing with lysis buffer, the protein was eluted with a linear gradient of 0-500 mM imidazole in 10 column volumes. The fractions of interest were pooled, concentrated with a 10 kDa cut-off Amicon concentrator (Millipore) and loaded onto a HiLoad Superdex 75 16/60 (GE Healthcare) equilibrated in SEC buffer (50 mM Sodium Phosphate buffer pH 7.0, 200 mM NaCl, 5% (w/v) glycerol and 1 mM TCEP).

Expression and purification of Bub1 and BubR1 constructs, as well as of Bub1:Bub3 and BubR1:Bub3 complexes was carried out as described (Breit et al., 2015; Overlack et al., 2015).

### Analytical SEC analysis

4 µM Bub1 and BubR1 protein constructs or Bub1:Bub3 and BubR1:Bub3 complexes were mixed with 16 µM CENP-E and CENP-F proteins respectively, in 30 µl final volume. Analytical size exclusion chromatography was carried out at 4°C on a Superose 6 5/150 or Superdex 75 5/150 in a buffer containing 50 mM HEPES pH 8.0, 100 mM NaCl, 5% (w/v) glycerol and 0.5 mM TCEP at a flow rate of 0.12 ml/min on an ÄKTA micro system. Elution of proteins was monitored at 280 nm,488 nm (eGFP-tag) and 587 nm (mCherry-tag). 50 µl fractions were collected and analysed by SDS-PAGE and Coomassie blue staining.

### Analytical Ultracentrifugation

Sedimentation velocity AUC was performed at 42,000 rpm at 20°C in a Beckman XL-A ultracentrifuge. Protein samples were loaded into standard double-sector centerpieces. The cells were scanned every minute and 500 scans were recorded for every sample. 6 µM mCherry-CENP-F^2866-2990^, mCherry-CENP-F^2922^- ^2990^ and mCherry-CENP-F^2950-2990^ were scanned at 587 nm. 7 µM eGFP-CENP-E^2070-C^ alone or mixed with 21 µM BubR1:Bub3 were instead scanned at 488 nm. Data were analyzed using the program SEDFIT (Brown and Schuck, 2006) with the model of continuous c(s) distribution. The partial specific volumes of the proteins, buffer density, and buffer viscosity were estimated using the program SEDNTERP. Data figures were generated using the program GUSSI (Brautigam, 2015).

### Protein Electroporation

For eGFP-CENP-E protein electroporation, HeLa cells were arrested in G2 with a 9 µM RO-3306 treatment for 15 hours (Millipore) and then released into mitosis for 3 hours in presence of 3.3 µM nocodazole. Mitotic cells were then collected by mitotic shake-off, washed with PBS and counted. Approximatively 3×10^6^ cells were then electroporated (Neon Transfection System Kit, Thermo Fisher) with 10 µM eGFP-CENP-E. Following electroporation, cells were allowed to recover in media with 3.3 µM nocodazole for 4 hours and then fixed and prepared for immunofluorescence analysis. For mCherry-CENP-F protein electroporation, HeLa cells were treated for 16 hours with 0.33 µM Nocodazole (Sigma) to synchronise cells in mitosis. Mitotic cells were then collected by mitotic shake-off, washed with PBS and counted. Approximatively 3×10^6^ cells were then electroporated with 5 µM mCherry-CENP-F. Following electroporation, cells were allowed to recover in media with 3.3 µM nocodazole for 4 hours and then fixed and prepared for immunofluorescence analysis.

### Low-angle metal shadowing and electron microscopy

mCherry-CENP-F^2688-C^ fractions from the elution peak of an analytical size-exclusion chromatography column were diluted 1:1 with spraying buffer (200 mM ammonium acetate and 60% glycerol) and air-sprayed as described (Baschong and Aebi, 2006; Huis in ‘t Veld et al., 2016) onto freshly cleaved mica pieces of approximately 2×3 mm (V1 quality, Plano GmbH). Specimens were mounted and dried in a MED020 high-vacuum metal coater (Bal-tec). A Platinum layer of approximately 1 nm and a 7 nm Carbon support layer were evaporated subsequently onto the rotating specimen at angles of 6-7° and 45° respectively. Pt/C replicas were released from the mica on water, captured by freshly glow-discharged 400- mesh Pd/Cu grids (Plano GmbH), and visualized using a LaB6 equipped JEM-1400 transmission electron microscope (JEOL) operated at 120 kV. Images were recorded at a nominal magnification of 60,000x on a 4k x 4k CCD camera F416 (TVIPS), resulting in 0.18 nm per pixel.

### Mammalian Plasmids

Plasmids were derived from the pCDNA5/FRT/TO-EGFP-IRES, a previously modified version (Krenn et al., 2012) of the pCDNA5/FRT/TO vector (Invitrogen). To create N-terminally-tagged EGFP-Bub1 truncation constructs, the Bub1 sequence was obtained by PCR amplification from the previously generated pCDNA5/FRT/TO-EGFP-Bub1-IRES vector (Krenn et al., 2012) and subcloned in frame with the GFP-tag. All Bub1 constructs were RNAi resistant (Kiyomitsu et al., 2007). pCDNA5/FRT/TO-based plasmids were used for generation of stable cell lines. All plasmids were verified by sequencing.

### Cell culture and transfection

HeLa cells were grown in DMEM (PAN Biotech) supplemented with 10 % FBS (Clontech), penicillin and streptomycin (GIBCO) and 2 mM L-glutamine (PAN Biotech). Flp-In T-REx HeLa cells used to generate stable doxycycline-inducible cell lines were a gift from S.S. Taylor (University of Manchester, Manchester, England, UK). Flp-In T-REx host cell lines were maintained in DMEM with 10 % tetracycline-free FBS (Clontech) supplemented with 50 µg/ml Zeocin (Invitrogen). Flp-In T-REx HeLa expression cell lines were generated as previously described (Krenn et al., 2012). Briefly, Flp-In T-Rex HeLa host cells were cotransfected with a ratio of 9:1 (w/w) pOG44:pcDNA5/FRT/TO expression plasmid using X- tremeGene transfection agent (Roche). 48 h after transfection, Flp-In T-Rex HeLa expression cell lines were put under selection for two weeks in DMEM with 10 % tetracycline-free FBS (Invitrogen) supplemented with 250 µg/ml Hygromycin (Roche) and 5 µg/ml Blasticidin (ICN Chemicals). The resulting foci were pooled and tested for expression. Gene expression was induced by addition of 0.5 µg/ml doxycycline (Sigma) for 24 h.

siBUB1 (Dharmacon, 5’-GGUUGCCAACACAAGUUCU-3’) or siBUBR1 (Dharmacon, 5’-CGGGCAUUUGAAUAUGAAA-3’) duplexes were transfected with Lipofectamine 2000 (Invitrogen) at 50 nM for 24 h. siCENP-E (Dharmacon, 5’- AAGGCUACAAUGGUACUAUAU-3’) and siCENP-F (Dharmacon, 5’- CAAAGACCGGUGUUACCAAG-3’ and 5’-AAGAGAAGACCCCAAGUCAUC-3’) duplexes were transfected at 60 nM with LipofectamineRNAiMAX (Invitrogen) for 24 h. siZwilch (SMART pool from Dharmacon, #L-019377-00-0005) duplexes were transfected with LipofectamineRNAiMAX at 120 nM for 72 h.

### Immunoblotting

To generate mitotic populations for immunoblotting experiments, cells were treated with 330 nM nocodazole for 16 h. Mitotic cells were then harvested by shake off and lysed in lysis buffer [150 mM KCl, 75 mM Hepes, pH 7.5, 1.5 mM EGTA, 1.5 mM MgCl_2_, 10 % glycerol, and 0.5 % Triton-X 100 supplemented with protease inhibitor cocktail (Serva) and PhosSTOP phosphatase inhibitors (Roche)]. Cleared cell lysates were resuspended in sample buffer, boiled and analyzed by SDS-PAGE using 3- 8 % gradient gels (NuPAGE^®^ Tris-Acetate Gels, Life technologies) and Western blotting. The following antibodies were used: anti-CENP-E (rabbit, ab133583, 1:500), anti-CENP-F (rabbit, Novus NB500-101, 1:500) and anti-Tubulin (mouse, Sigma T9026, 1:10000). Secondary antibodies were anti–mouse (Amersham) or anti–rabbit (Amersham) affinity-purified with horseradish peroxidase conjugate (working dilution 1:10000). After incubation with ECL Western blotting system (GE Healthcare), images were acquired with the ChemiDoc^™^MP Imaging System (BioRad) in 16-bit TIFF format. Images were cropped and converted to 8-bit using Image J software (NIH). Brightness and contrast were adjusted using Photoshop CS5 (Adobe).

### Live cell imaging

Cells were plated on a 24-well µ-Plate (Ibidi®). The medium was changed to CO_2_ Independent Medium (Gibco®) 6 h before filming. DNA was stained by addition of the SiR-Hoechst-647 Dye (Spirochrome) to the medium 1 h before imaging. Cells were imaged every 5 to 10 min in a heated chamber (37 °C) on a 3i Marianas(tm) system (Intelligent Imaging Innovations Inc.) equipped with Axio Observer Z1 microscope (Zeiss), Plan-Apochromat 40x/1.4NA oil objective, M27 with DIC III Prism (Zeiss), Orca Flash 4.0 sCMOS Camera (Hamamatsu) and controlled by Slidebook Software 6.0 (Intelligent Imaging Innovations Inc).

### Immunofluorescence

HeLa cells and Flp-In T-REx HeLa cells were grown on coverslips precoated with poly-D-Lysine (Millipore, 15 µg/ml) and poly-L-Lysine (Sigma), respectively. Asynchronously growing cells or cells that were arrested in prometaphase by the addition of nocodazole (Sigma-Aldrich) were fixed using 4 % paraformaldehyde. Cells were stained for Bub1 (mouse, ab54893, 1:400), BubR1 (rabbit, Bethyl A300-386A-1, 1:1000), Tubulin (mouse, DM1a Sigma, 1:500), CENP-E (mouse, ab5093, 1:200), CENP-F (rabbit, Novus NB500-101, 1:300), Zwilch (rabbit, in-house made, SI520, 1:900), Mad1 labelled with AlexaFluor-488 (mouse, in-house made, Clone BB3-8, 1:200), pT232-AurB (rabbit, Rockland #660-401-667, 1:2000), Plk1 (mouse, ab17057, 1:300), pS10H3 (mouse, ab14955, 1:3000), pT121 H2A (rabbit, active motif #39391, 1:2000) and CREST/anti-centromere antibodies (Antibodies, Inc., 1:100), diluted in 2 % BSA-PBS for 1.5 h.

For testing the effect of various kinase inhibitors on CENP-E and CENP-F kinetochore localization, the protocol was adapted in the following way: Cells were pre-permeabilized with 0.5 % triton-X-100 solution in PHEM (Pipes, Hepes, EGTA, MgCl2) buffer for 2 min before fixation with 4% PFA-PHEM for 15 min. After blocking the cells with 3 % BSA-PHEM buffer supplemented with 0.1 % triton-X-100, they were incubated at room temperature for 1-2 h with primary antibodies diluted in blocking buffer. Washing steps were performed in PHEM-T buffer.

Goat anti–human (Invitrogen), goat anti–mouse (Jackson ImmunoResearch Laboratories, Inc.) and goat anti-rabbit (Jackson ImmunoResearch Laboratories, Inc.) fluorescently labeled antibodies were used as secondary antibodies. DNA was stained with 0.5 µg/ml DAPI (Serva) and coverslips were mounted with Mowiol mounting media (Calbiochem). Cells were imaged at room temperature using a spinning disk confocal device on the 3i Marianas(tm) system equipped with an Axio Observer Z1 microscope (Zeiss), a CSU-X1 confocal scanner unit (Yokogawa Electric Corporation), Plan-Apochromat 63x or 100x/1.4NA Oil Objectives (Zeiss) and Orca Flash 4.0 sCMOS Camera (Hamamatsu). Images were acquired as z-sections at 0.27 µm. Images were converted into maximal intensity projections, exported and converted into 8-bit. Quantification of kinetochore signals was performed on unmodified 16-bit z-series images using Imaris 7.3.4 32-bit software (Bitplane). After background subtraction, all signals were normalized to CREST. At least 117 kinetochores were analyzed per condition. Measurements were exported in Excel (Microsoft) and graphed with GraphPad Prism 6.0 (GraphPad Software).

### Cell synchronization

To test the effect of various kinase activities on CENP-E and CENP-F kinetochore localization, cells were synchronized using a double thymidine arrest. Cells were released from the first 18 h thymidine (2 mM; Sigma–Aldrich) block by washing them with fresh pre-warmed media several times. After releasing them for the next 9 h, cells were exposed to thymidine (2 mM) a second time for 15 h. Afterwards, cells were released into S-phase for 4 h and then nocodazole (330 nM) was added to the media for the next 3-4 h to enrich for the mitotic cell population. Kinase activity inhibitors, BI 2536 (500 nM; Calbiochem), Hesperadin (500 nM; Calbiochem), Reversine (500 nM; Calbiochem) or BAY-320 (10 µM; kindly received from Dr. Gerhard Siemeister, Bayer GmbH, Berlin) were added in the presence of the proteasome inhibitor, MG132 (10 µM; Calbiochem) to the cells for 90 min before fixing these cells for immunofluorescence.

### Chromosome alignment

For analysis of the effect of CENP-F depletion on chromosome alignment, cells were fixed after RNAi either asynchronously or after an additional treatment with 10 µM MG-132 for 2 h. Cells were stained for CENP-F, Tubulin and CREST. DNA was labeled with DAPI. The number of metaphase cells with aligned chromosomes and with misaligned chromosomes was scored for each condition. At least 595 cells (without synchronization) or 92 cells (with synchronization) were analyzed per condition.

